# CENO: A Genome-Scale World Model for Evolutionary Sequence Interpretation and Programmable Regulatory Design

**DOI:** 10.64898/2026.07.28.741284

**Authors:** Mingqian Ma, Yucheng Wu, Xin Chen, Feifei Jiang, Peijun Lin, Dongxin Ye, Yidi Sun, Yijing Zhang, Tianqiong Shi, Yu Zhao, Wanli Ouyang, Bowen Zhou, Lei Bai, Yuchen Ren

**Affiliations:** Shanghai Artificial Intelligence Laboratory; Fudan University; The University of Sydney; Shanghai Jiao Tong University; University of Southern California; Shanghai Innovation Institute; University of Electronic Science and Technology of China; Center for Excellence in Brain Science and Intelligence Technology Chinese Academy of Sciences; Nanjing Normal University; Westlake University; Shenzhen Loop Area Institute; The Chinese University of Hong Kong; Tsinghua University

## Abstract

DNA encodes biological function across a continuum of sequence scales, from single-nucleotide and motif-level grammar to regulatory neighborhoods, chromatin-scale organization and evolutionary constraint. A useful model of genomes should therefore do more than classify short sequence windows: it should maintain nucleotide-resolution state over long contexts, score counterfactual mutations, condition on homologous sequence evidence and generate candidates that can be evaluated against structural or functional objectives. We define such a system operationally as a genomic world model: a general-purpose generative model of genome sequence space that unifies sequence understanding and sequence design through a shared state and likelihood interface. Here we introduce CENO, a family of long-context generative genomic world models designed to preserve local DNA grammar while extending usable context to regulatory and chromatin scales. CENO combines Mamba sequence-mixing layers, sparse attention layers and mixture-of-experts capacity in a single autoregressive backbone, and is trained at 300M, 600M and 1B parameter scales with a staged curriculum that progresses from 8k-token cross-domain genomic pretraining to 131k- and 1M-token whole-genome long-context continuation. We evaluate CENO under a unified world-model benchmark paradigm spanning retrieval, representation, counterfactual perturbation, reconstruction, evolutionary conditioning and design. CENO retains practical long-context inference and retrieves distal sequence in synthetic assays. In zero-shot long-context analyses, without task-specific fine-tuning, long-context continuation yields annotation- and chromatin-boundary-associated attention patterns and frozen-state representations that generalize across human cell types and mouse cell or tissue settings. To incorporate evolutionary information, we further post-train CENO on packed real multiple-sequence-alignment contexts and score variants by reference–mutant likelihood deltas, improving matched variant-effect prediction and producing evolutionary enrichment signals across species. Complementing these perturbation-based variant tests, we evaluate zero-shot long-sequence generation by partial-gene continuation, asking whether the model can recover withheld gene-scale sequence structure across eukaryotic, bacterial and archaeal species; recovery improves with model scale and later whole-genome long-context training. Finally, we use CENO as the backbone for a cell-type-specific enhancer design workflow in mouse cortex, coupling a CENO-based accessibility oracle with conditional supervised fine-tuning and oracle-guided reinforcement learning. Together, CENO provides a genome-scale sequence world-model framework for sequence interpretation, evolutionary reasoning, gene-scale reconstruction and programmable regulatory sequence generation.

## 1. Introduction

Genome modeling is a short-to-long sequence-learning problem. At short ranges, DNA encodes nucleotide-resolution syntax, transcription-factor motifs, splice signals, codon structure and local regulatory grammar over tens to hundreds of bases. At longer ranges, the same sequence is organized into promoters, enhancers, repeats, genes, regulatory neighborhoods, chromatin domains and locus-scale structures that can span tens of kilobases to megabases. A useful genomic model should therefore not trade local precision for long context, or long context for local grammar. It should connect these scales in a single sequence model, learning how short sequence features are composed, routed and constrained across progressively longer genomic contexts.

We use the term genomic world model in an operational sense. The “world” is not a complete organismal or cellular simulator, but the sequence-level world in which local nucleotide grammar, gene structure, regulatory neighborhoods, chromatin-scale organization, evolutionary constraint and designable function jointly determine the plausibility and consequence of DNA sequences. Under this definition, a genomic world model should satisfy four requirements. First, it should be general-purpose, spanning coding, noncoding, regulatory and cross-species sequence regimes rather than a single supervised task. Second, it should be stateful over long contexts, so that distal sequence can influence local predictions and internal representations reflect locus-scale organization. Third, it should unify understanding and generation: the same model should score mutations, recover withheld sequence, condition on homologous context and generate new sequences. Fourth, it should be evaluated by a unified benchmark suite that probes retrieval, representation, perturbation, reconstruction, evolution and design rather than by a single leaderboard. In this sense, a genomic foundation model is defined by reuse, whereas a genomic world model is defined by reusable operations over genome sequence space: completion, perturbation, conditioning and design through a shared generative interface.

Recent DNA language models have advanced different parts of this goal. Encoder-style models such as DNABERT-2, Nucleotide Transformer and Genomics-FM showed that large-scale pretraining can produce transferable genomic representations [1, 2, 3]. Long-sequence architectures such as HyenaDNA and Caduceus extended genomic modeling beyond the context lengths of standard Transformers [4, 5]. Generative models including Evo, Evo 2, HybriDNA and OmniReg-GPT further moved the field toward autoregressive genome-scale sequence modeling and sequence generation [6, 7, 8, 9]. In parallel, supervised sequence-to-function models such as Enformer, Borzoi and AlphaGenome have shown that DNA sequence models can predict regulatory activity, expression and variant effects when trained directly on molecular phenotypes [10, 11, 12]. Together, these studies show that DNA models can predict local sequence grammar, long-range dependencies, molecular phenotypes and generative sequence distributions. However, these capabilities are often developed and evaluated in separate modeling regimes. Existing genome-scale generative models have largely focused on scaling sequence modeling itself; whether long-context continuation produces reusable genomic representations, evolutionary reasoning and controllable design remains less explored.

This separation leaves an important gap for genomic world modeling. Long context is not equivalent to a large maximum input length. A model may accept long sequences while still failing to retrieve distant information, generate efficiently after long prompts, or form internal representations that reflect long-range genomic organization. Conversely, a model optimized for local or short-window tasks may perform well on motif, splicing or coding readouts while remaining insensitive to regulatory context beyond the immediate neighborhood. A practical long-context genomic world model should therefore satisfy three conditions: it should remain computationally usable at long input lengths; it should preserve short-range sequence grammar while routing information across distant loci; and it should expose representations that can be interrogated for biologically meaningful long-range structure.

Benchmark behavior also reflects this short-to-long tension. Performance on short regulatory classification, splice-altering variants, coding fitness, structural variants, long-context retrieval, chromatin-scale diagnostics and sequence generation can favor different model families. Variant-effect prediction and sequence generation probe complementary aspects of genomic understanding. Variant-effect prediction tests whether local perturbations receive meaningful scores within an observed genomic context. Generation asks both whether a model can reconstruct withheld natural sequence and whether its generative distribution contains candidates for new biological objectives. Together, these tasks test whether model scaling and long-context training improve both interpretation and guided exploration of sequence space. A genomic world model should therefore be evaluated not only by variant scores, but also by whether its generative distribution improves with scale and whole-genome long-context training.

Generalization across biological contexts is another requirement. A model that detects a chromatinboundary-associated signal in one locus or one cell type may not have learned a reusable long-range representation. Similarly, a model that recovers sequence continuations in one species may not have learned a broadly useful genomic distribution. Long-context evaluation should therefore test whether model-internal signals transfer across cell types, tissues and species, and whether sequence recovery holds across diverse genomes rather than only within a single reference context. This generalization requirement is central to the world-model view: the learned sequence distribution should support operations that transfer across biological contexts, not only memorized local patterns.

Variant interpretation adds a further axis to this problem. Many variant effects are shaped not only by the target sequence itself, but also by evolutionary constraint across homologous sequences. Alignment- and phylogeny-aware models such as GPN-MSA and GPN-Star use multispecies genome alignments to improve variant-effect prediction [13, 14]. PoET, developed for protein families, further shows that family-level evolutionary context can provide a strong training signal for autoregressive sequence models [15]. These studies point to a missing capability for genomic world models: combining single-sequence genome pretraining with post-training that exposes the model to homologous context, while retaining a simple sequence-likelihood interface for downstream variant scoring.

A further goal is to move from interpretation to design. Genomic foundation models are useful not only when they score existing variants or annotate existing loci, but also when they can generate candidate sequences under functional constraints. Regulatory design makes the short-to-long problem especially clear. A designed enhancer must contain local motif grammar, preserve plausible sequence composition, and satisfy cell-type-specific functional constraints defined by a broader regulatory program. This requires a generative backbone that can be adapted with task-specific feedback without discarding the sequence grammar and contextual representations learned during pretraining. In the world-model framing, design is not a separate downstream add-on: it is one of the core operations that tests whether the learned sequence distribution can be steered toward new functional regions of genome space.

Here we introduce CENO, a family of long-context generative genomic world models designed around this short-to-long principle. CENO combines Mamba sequence-mixing layers, sparse attention layers and mixture-of-experts layers in a single causal autoregressive backbone. We trained 300M-, 600M- and 1B-parameter models with a staged curriculum. The first stages learned broad genomic sequence grammar with an 8k-token context across prokaryotic, metagenomic, viral, organelle, eukaryotic, transcript, splicing and regulatory sources. The later stages shifted toward eukaryotic, transcript, splicing and regulatory sequence and then extended the context to 131k and 1M tokens using long windows from complete eukaryotic genomes. This curriculum separates three axes that are often entangled in DNA language modeling: model capacity, sequence length and data composition. It also creates a natural test of whether later whole-genome continuation improves gene-scale sequence recovery, rather than only increasing the configured context length.

CENO is also designed for evolutionary post-training. The pretrained model first learns single-sequence genome syntax and long-range state tracking. We then adapt the same backbone to packed real multiple-sequence-alignment contexts, in which a target sequence is paired with homologous rows and explicit row identities. Early row-local layers preserve within-sequence modeling, whereas later fusion layers allow homologous rows to condition the target representation. At inference, variant effects are computed as likelihood differences between mutant and reference target sequences under the same MSA context. Thus, evolutionary information shapes the model during post-training and scoring, while the downstream readout remains a direct sequence-likelihood delta rather than a supervised task-specific classifier.

We evaluate CENO across the capabilities required of a genomic world model. We first measure pretraining scaling, long-prefill throughput and sustained generation speed, and test whether distant sequence influences terminal predictions in a synthetic DNA needle-in-a-haystack assay. We then probe whether long-context continuation induces annotation- and chromatin-boundary-associated signals, using zero-shot TAD-scale attention diagnostics, annotation-linked attention examples and frozen-state linear probes across human cell types and mouse cell or tissue settings. We next evaluate zero-shot genomic prediction across variant-effect, fitness, regulatory-impact and structural-variant benchmarks. Complementing these perturbation-based tests, we evaluate long-sequence generation by partial-gene continuation across eukaryotic, bacterial and archaeal species, treating withheld-sequence recovery as a reconstruction-style benchmark for gene-scale sequence understanding. We further test whether CENO can supply long insertion candidates for structure-guided design across two boundary-deletion loci and a duplication-induced neo-domain. Candidate sequences are selected using one predictive structure model and evaluated using a second model for boundary-associated contact recovery or strengthened insulation.

We then evaluate MSA-context post-training for evolutionary variant scoring across matched BRCA1/BRCA2 comparisons, human disease and fine-mapping benchmarks, and evolutionary enrichment tasks across species. Finally, we use CENO as the backbone for a cell-type-specific enhancer design workflow in mouse cortex, coupling a CENO-based accessibility oracle with conditional supervised fine-tuning and GRPO-based reinforcement learning. Together, these analyses position CENO as a genome-scale world-model framework that links local sequence grammar, long-range genome organization, counterfactual variant scoring, evolutionary conditioning, gene-scale sequence recovery and regulatory sequence generation.

## 2. Results

### 2.1. CENO pretraining spans local genome grammar to megabase context

CENO was trained as a family of causal genomic language models with approximately 300M, 600M and 1B total parameters. Figure 1a outlines a long-context pretraining program that links the model backbone, context curriculum and downstream interfaces. This design provides a shared foundation for using extended genomic context in analysis, downstream applications and sequence generation.

**Figure 1.**
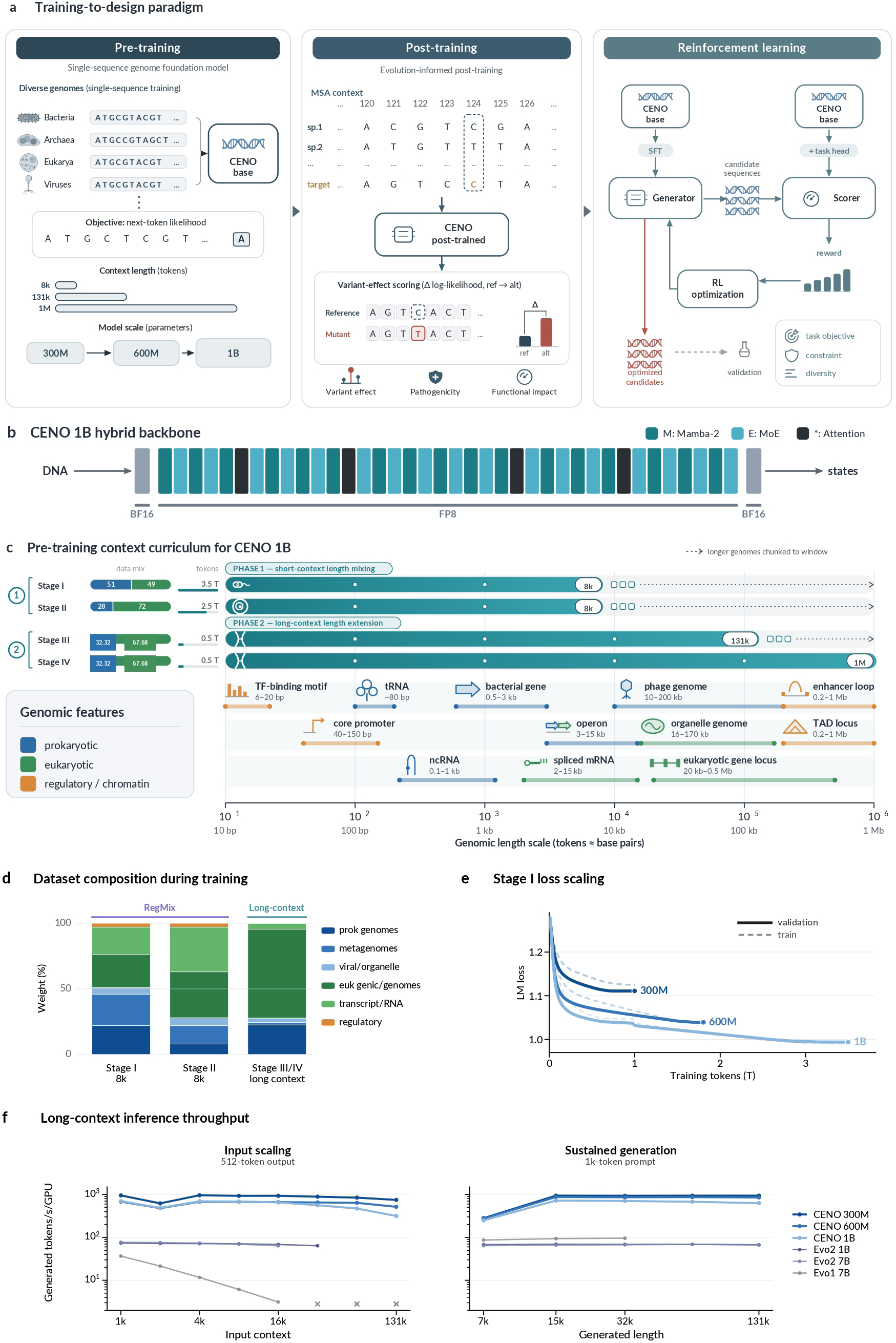
Overview of the CENO genomic foundation model. **a,** Training-to-design paradigm. *Pretraining*: a single-sequence next-token-likelihood objective over diverse genomes (bacteria, archaea, eukarya, viruses) yields the CENO base model under a context-length curriculum (8k → 131k → 1M tokens) at three scales (300M/600M/1B). *Post-training*: real multiple-sequence alignments (MSA) supply evolutionary constraint; the post-trained model gives a variant-effect readout (VEP score) from the log-likelihood difference (Δ log *p*) between reference and alternate alleles, supporting VEP, pathogenicity and functional-impact tasks. *Reinforcement learning*: from the CENO base, supervised fine-tuning (SFT) and a task head yield a CENO generator and scorer; candidate sequences are generated and scored, and the reward drives RL optimization toward higher-reward candidates under task objectives, constraints and diversity goals (dashed arrow, validation). Color encodes input/data, model, evaluation and output roles. **b,** CENO 1B hybrid backbone: a causal trunk interleaves Mamba-2 (M), mixture-of-experts (E) and attention (∗) blocks between the input and output embeddings; embeddings use BF16, and trunk computation uses FP8. **c,** Biological context-length curriculum on a shared log length axis (tokens ≈ base pairs). Top: every stage trains on the full length range, with a solid bar for the contiguous context window and a segmented bar for longer genomes chunked to fit it (Stage I–II, 8k; Stage III, 131k; Stage IV, 1M). Bottom: representative genomic features at their characteristic length. **d,** Dataset composition during training: stacked weight (%) per stage; Stage I/II set by constrained RegMix, Stage III/IV by long-context weighting. **e,** Stage I loss scaling: validation (solid) and train (dashed) LM loss versus training tokens for 300M/600M/1B; the 1B model reaches the lowest endpoint. **f,** Long-context inference throughput (tokens s^−1^ GPU^−1^): input scaling with 512-token output (left) and sustained generation from a 1k-token prompt (right), for CENO 300M/600M/1B versus Evo 2 1B/7B and Evo 1 7B; crosses (×) mark runs that did not complete (out of memory). Image created with BioRender.com, with permission.

The backbone combines Mamba-2 sequence-mixing blocks, attention blocks and mixture-of-experts (MoE) capacity within a single causal architecture (Figure 1b, Methods 4.1). Across the 300M, 600M and 1B total parameter scales, the corresponding active parameter counts were approximately 100M, 200M and 400M per token.

The context curriculum trained Stage I/II models at 8,192 tokens. Stage III continued from the 8k checkpoint at 131,072 tokens. Stage IV continued from the 131k checkpoint at 1,048,576 tokens (Figure 1c). Stage I/II production recipes were selected by a constrained RegMix workflow (Methods 4.2). Candidate mixtures were sampled under eukaryotic/prokaryotic balance constraints, fitted from small-scale mixture-response validation experiments and selected with prespecified benchmark-response filters. The fixed production weights are reported in Supplementary Table S2 and source data. The additional selection archives needed to recreate the mixture search are identified separately. At the high-level category scale, this produced a near-balanced Stage I recipe: 51% prokaryotic, metagenomic, viral and organelle data, and 49% eukaryotic, transcript, splicing and regulatory data. Stage II shifted this grouped split to 28%/72%. Stage III/IV then used separate long-context weighting with a 32.32%/67.68% grouped split. These long-context stages added complete eukaryotic-genome sources sampled as long training windows (Figure 1d).

Within the shared Stage I 8k regime, loss curves followed model scale. Training and validation losses decreased across consumed tokens for all three displayed model sizes. The 1B model reached the lowest endpoint validation loss among the 300M, 600M and 1B runs (Figure 1e). These curves isolate model-size scaling within a fixed context regime.

CENO retained practical throughput at long context. In the long-prefill benchmark, output length was fixed at 512 tokens while input length increased. At 131,072 input tokens, the CENO 300M, 600M and 1B models reached 747.41, 515.83 and 315.10 generated tokens per second per GPU, respectively. Two Evo 2 baselines did not complete this input length in the same setup because of out-of-memory errors (Figure 1f; Methods 4.3). Failed comparator runs at requested lengths are retained in the source data and categorized by failure class. In sustained long-context decoding, a 1,024-token prompt was extended to 131,072 total tokens. At the terminal point, CENO 300M, 600M and 1B reached 938.97, 852.03 and 627.57 generated tokens per second per GPU, respectively. Evo 2 1B and Evo 2 7B reached 66.87 and 67.38 generated tokens per second per GPU. Together, these results establish CENO as a practical long-context backbone, supporting stronger genomic analysis, downstream applications and sequence generation.

### 2.2. Long-context CENO links distant sequence to chromatin-scale readouts

Genomic language models can now be trained on long sequence windows, but whether they use distant context in biologically relevant tasks remains a central question. Having established the CENO backbone, we tested long-context use with readouts that move from controlled sequence retrieval to chromatin-scale structure. A synthetic needle assay asked whether distant bases affected prediction. Topologically associating domain (TAD)-scale attention and annotation-linked examples tested whether long-context signals aligned with chromatin organization and regulatory elements. Frozen-state probes then asked whether boundary information was encoded in hidden representations.

We first asked whether CENO could use sequence placed far from the prediction site. In the DNA needle-in-a-haystack assay, a 100-bp sequence was inserted into a random DNA background at 10%, 50% or 90% depth and repeated at the terminal query position. Mutating the inserted needle changed the A/T/C/G log-probability vector at the corresponding query base. The retrieval score summarized this perturbation across the 100 needle positions (Methods 4.4).

The CENO 300M, 600M and 1B checkpoints retained retrieval scores above threshold across the 32k, 64k, 131k and 1M contexts shown in Figure 2a. Retrieval generally improved with model size and later training stage, particularly at longer contexts. This assay showed that distant sequence can affect terminal predictions.

**Figure 2.**
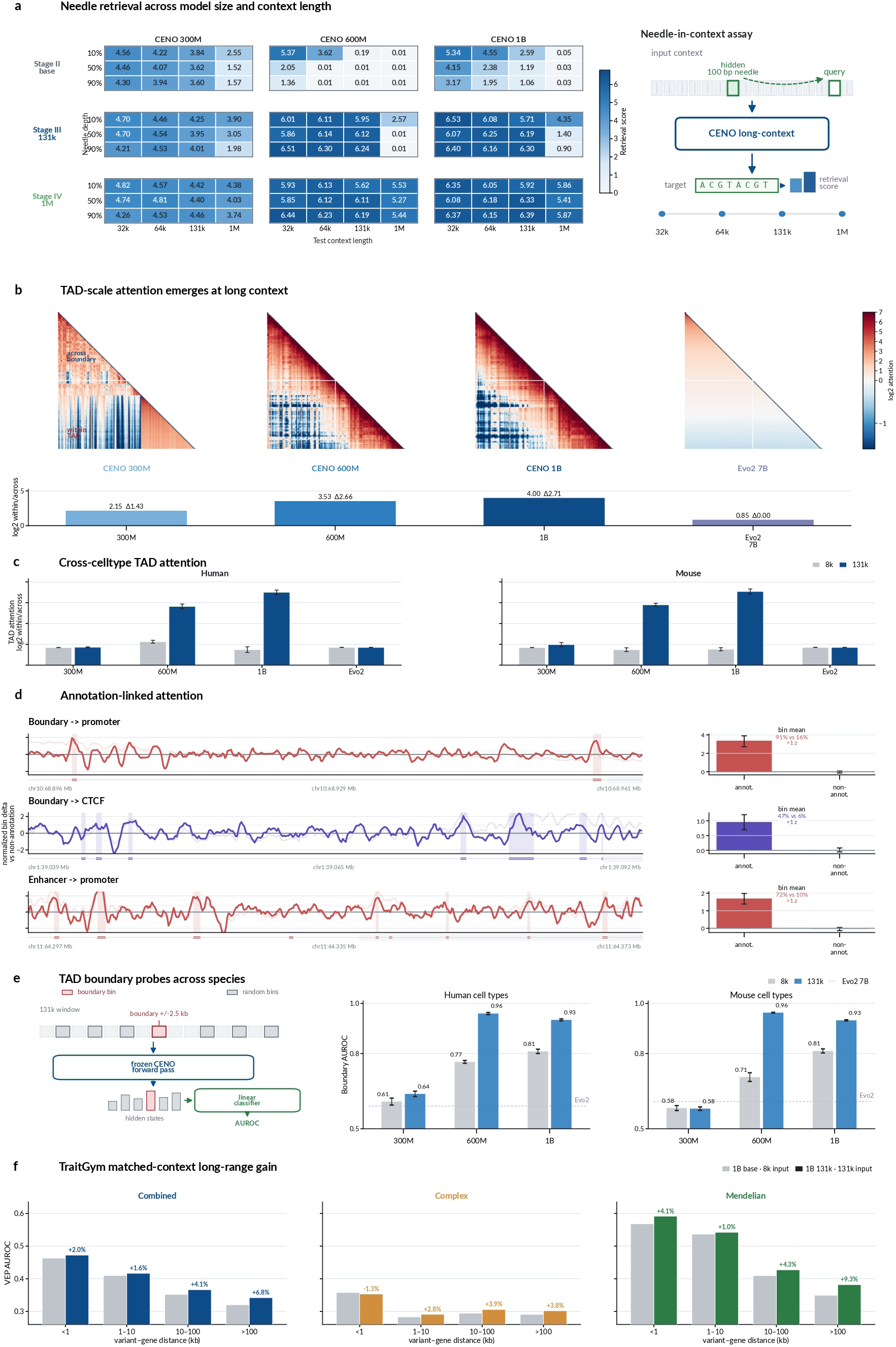
Long-context retrieval and chromatin-scale readouts. **a,** DNA needle retrieval across 32k, 64k and 131k contexts. **b,** Matched-locus TAD attention maps and 131k/8k within/across-boundary gain; the 24-locus audit and retained examples are in the Supplementary material. **c,** Cross-cell-type TAD attention summaries across five human cell types and five mouse cell or tissue settings. **d,** Selection-audited annotation-linked attention examples, bounded to locus-level support. **e,** Frozen-state TAD boundary probes across human and mouse settings; selected-layer, cell-level spread and label-shuffle controls are retained in source data. **f,** TraitGym matched-context long-range gain across three trait groups (Combined, Complex, Mendelian): each sub-panel pairs the 1B base scored at 8k input (gray) against the 1B Stage III model scored at 131k input (color) by variant–gene distance, with the relative gain shown above each bin; the benefit grows at longer range and is largest for Mendelian variants beyond 100 kb.

We next asked whether model attention, without supervised boundary labels, associated with chromatin-domain organization. TADs provide a direct readout because loci on the same side of a boundary are expected to interact more strongly than loci separated by that boundary. Because both CENO and Evo 2 contain attention layers, we directly compared their attention maps at matched TAD-boundary loci. Maps were scored as log_2_ within-boundary attention relative to across-boundary attention. Context scaling was assessed by comparing matched Stage II 8k and 131k windows at the same locus. In the representative locus shown in Figure 2b, 131k continuation increased this within/across-boundary score for CENO 300M, 600M and 1B by 1.43, 2.66 and 2.71 log_2_ units, respectively. By contrast, the Evo 2 7B reference remained essentially unchanged in the same locus. We then tested whether the TAD attention signal generalized beyond one displayed locus and one cell context. The cross-setting analysis covered five human cell types and five mouse cell or tissue settings, with three CENO model scales and an Evo 2 7B reference evaluated under matched 8k and 131k contexts. Across the five human cell types, mean log_2_ within/across-boundary attention increased from 1.13 to 2.82 for CENO 600M after 131k continuation. For CENO 1B, the same score increased from 0.74 to 3.50 (Figure 2c, Methods 4.5). Across five mouse cell or tissue settings, the corresponding values increased from 0.74 to 2.90 for CENO 600M and from 0.76 to 3.54 for CENO 1B. By contrast, CENO 300M and Evo 2 7B remained near baseline. Thus, the 131k continuation effect was not restricted to the representative locus, but generalized across cell-type and species settings in the 600M and 1B CENO models.

We next inspected whether high-attention regions overlapped known regulatory annotations in selected loci. This analysis was also performed zero-shot, without explicit annotation supervision. We show three CENO 1B examples: boundary-to-promoter, boundary-to-CTCF-bound candidate cisregulatory element (cCRE) and enhancer-to-promoter attention. In these examples, annotated bins had higher normalized attention than non-annotated bins. The fractions above one normalized unit were 91% versus 16%, 47% versus 6% and 72% versus 10%, respectively (Figure 2d; Methods 4.6). These examples show that CENO concentrates attention on functionally related sequence, linking chromatin-boundary readouts to regulatory annotations.

Finally, we tested whether chromatin-boundary information was encoded in hidden states of the model. A simple linear probe was trained to distinguish TAD boundary bins from matched random bins while the CENO backbone was held frozen. In human cell types, mean boundary AUROC increased from 0.767 at 8k to 0.960 at 131k for CENO 600M. For CENO 1B, the same metric increased from 0.809 at 8k to 0.934 at 131k (Figure 2e; Methods 4.7). In mouse cell or tissue settings, the corresponding increases were 0.706 to 0.963 for CENO 600M and 0.811 to 0.933 for CENO 1B. The 300M model showed only small changes in the same probe readout.

We next asked whether long-context pretraining translated into a downstream functional benefit on variant-effect prediction. On the TraitGym benchmark, we compared the 1B base model scored with an 8k-token input against the 1B Stage III model scored with a 131k-token input. Model context and evaluation input length therefore increased together. We stratified variants by variant–gene distance (Figure 2f; Methods 4.8). Matching longer context to longer input raised zero-shot AUROC in almost every distance bin, and the gain grew with distance: for the Combined trait set the relative improvement rose from +2.0% below 1 kb to +6.8% beyond 100 kb. The effect was strongest for Mendelian traits, reaching +9.3% beyond 100 kb, while Complex traits improved more modestly (+3.8% beyond 100 kb). Thus the additional context acquired during long-context pretraining yielded the largest functional gains precisely where the variant and its target gene were most distant.

Together, these assays show that long-context extension enables CENO to use distal sequence and encode chromatin-scale structure beyond one-dimensional sequence proximity.

### 2.3. CENO zero-shot scoring generalizes across variant-effect tasks

Identifying genetic variants that drive a biological function or disease phenotype remains a major challenge. Deep learning has shown considerable potential for addressing this problem. To evaluate sequence models systematically, we assembled a comprehensive variant-effect benchmark spanning diverse biological processes and functional consequences (Figure 3a).

**Figure 3.**
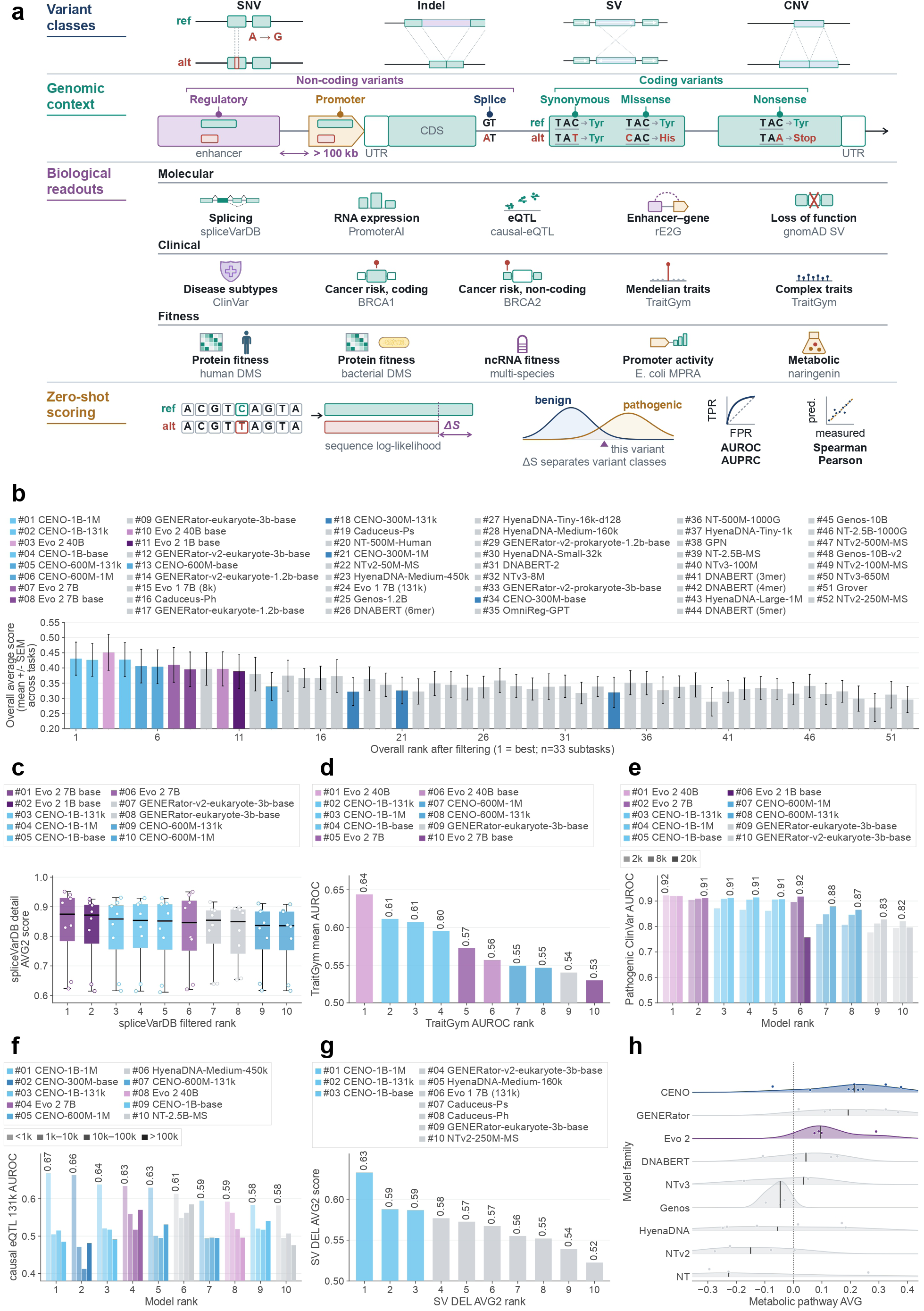
Cross-task zero-shot variant-effect benchmark. **a,** Benchmark scope and zero-shot scoring procedure. From top: variant classes, each drawn as the reference allele above and the derived allele below; genomic context, showing the positions a variant can occupy in and around a gene and how these divide into non-coding and coding consequences, with one reference codon (TAC, Tyr) carrying the synonymous, missense and nonsense outcomes; the fifteen biological readouts covered by the benchmark, grouped as molecular, clinical and fitness and labelled with their source datasets; and the scoring procedure, in which matched reference and alternate windows centred on the variant are scored by sequence likelihood and their difference Δ*S* is evaluated with classification (AUROC, AUPRC) or correlation (Spearman, Pearson) metrics without task-specific fitting. The fifteen readouts aggregate into the ten task families used for the overall ranking in **b**. **b,** Overall average score across the 33 selected subtasks, where evaluated, for all 52 models ordered by overall rank; markers show the mean and whiskers the s.e.m. across subtasks. Each subtask is taken at 8k input where a task offers several input lengths, so the ranking is a matched-context comparison. CENO checkpoints are highlighted. Three model families not discussed in the text (CENO-80M, CENO-7B and Carbon) are excluded from every panel. **c,** spliceVarDB AVG2 for the ten highest-ranked models; dots are the eight detail subtasks and the vertical rule the summary AVG2. **d,** TraitGym mean AUROC over 26 retained subtasks; AlphaGenome RNA-seq is set aside because it covers fewer than half of them. **e,** Pathogenic ClinVar AUROC at 2k, 8k and 20k input, with models ordered by the mean of the three. **f,** Causal eQTL AUROC at 131k input, stratified by variant-to-transcription start site (TSS) distance, with models ordered by the <1 kb stratum. **g,** Structural-variant deletion AVG2 for the ten highest-ranked models. **h,** Metabolic-pathway score by model family, for the nine families with at least three evaluated checkpoints; dots are individual checkpoints and the vertical rule the family median, with CENO checkpoints pooled across pre-training stages. AVG2 denotes the mean of paired AUROC and AUPRC scores where both metrics are available.

At the molecular level, the PromoterAI [16] and causal-eQTL datasets characterize the effects of genetic variants on RNA expression, whereas spliceVarDB [17] evaluates variant-induced alterations in RNA splicing. The structural-variant (SV) dataset [18] assesses whether large-scale genomic alterations lead to loss of function in specific genes. At the organismal and phenotypic levels, ClinVar [19], the BRCA1/2 [20, 21, 22] and TraitGym [23] datasets evaluate the disease-associated consequences of genetic variation. ClinVar supplies expert-reviewed pathogenicity labels across clinical categories, the BRCA1/2 datasets focus on variants in breast cancer susceptibility genes, and TraitGym evaluates variants associated with Mendelian and complex traits. In addition, we incorporated fitness datasets that provide an evolutionarily integrated measure of sequence functionality [24, 25, 26, 27, 28], together with 3^′^UTR, 5^′^UTR and mRNA-degradation reporter assays that probe post-transcriptional regulation, a long-range enhancer–gene interaction dataset [29] that captures regulatory relationships influencing gene expression, and a dataset in which different promoter–coding sequence combinations modulate naringenin production through their effects on metabolic pathways [30].

Beyond its broad coverage of biological functions, this benchmark is more comprehensive than conventional variant-effect datasets in three important respects. First, in addition to common single-nucleotide variants, the SV dataset includes a large collection of structural variants – deletions, insertions and copy-number gains – and their effects on gene function, and the spliceVarDB dataset incorporates insertion and deletion (indel) variants. Second, whereas many existing benchmarks primarily focus on coding regions, our collection contains extensive functional-effect data for noncoding regions, including ncRNA fitness, TraitGym and SV datasets. Third, the benchmark captures regulatory effects across multiple genomic scales. The enhancer–gene dataset includes enhancer– promoter interactions spanning more than 100 kb, while the causal-eQTL dataset contains long-range associations between regulatory variants and their target genes.

Overall, this benchmark encompasses diverse biological processes, variant classes, genomic contexts and interaction scales. It therefore provides a broad and comparatively balanced framework for evaluating how well genomic foundation models predict the functional consequences of sequence variation.

We next asked whether the pretrained CENO model family could score sequence perturbations without task-specific fitting [2, 13, 7, 12]. Task-specific predictors such as SpliceAI and Enformer are instead trained for one readout at a time [31, 10]. For each variant, we constructed matched reference and alternate windows and used the difference in sequence likelihood as a zero-shot effect score. We evaluated this score across ten benchmark families, which resolve into the fifteen biological readouts shown in Figure 3a: splice disruption, RNA expression and eQTL effects, enhancer–gene links, loss of function, clinical pathogenicity, cancer risk in coding and noncoding sequence, Mendelian and complex traits, protein and ncRNA fitness, promoter activity, and metabolic flux.

The aggregate benchmark revealed a consistent signal across model scales and training stages. Ranking all 52 evaluated models on their mean score over the 33 selected subtasks, taken at matched 8k input, CENO-1B-1M and CENO-1B-131k placed first and second, while the base checkpoint placed fourth (Figure 3b). The 131k and 1M CENO-600M checkpoints also ranked among the top six. Thus, the aggregate result reflected a family-wide pattern rather than an isolated best checkpoint. It did not, however, imply that CENO led every benchmark.

Task-level results exposed this heterogeneity. In spliceVarDB, Evo 2 7B base and Evo 2 1B base occupied the first two positions. The three CENO-1B checkpoints followed, with summary AVG2 scores spanning 0.824–0.828 (Figure 3c). TraitGym produced a different leaderboard. Evo 2 40B led, whereas the CENO-1B checkpoints formed the next cluster, with mean AUROC values spanning 0.595–0.611 (Figure 3d). CENO therefore transferred consistently, but other pretrained models retained an advantage on individual benchmarks.

Context length mattered most clearly for ClinVar. The 131k and 1M CENO-1B checkpoints gained approximately 0.04–0.05 AUROC when the input increased from 2k to 20k, ranking third and fourth behind Evo 2 40B and Evo 2 7B (Figure 3e). The effect was not universal. In causal eQTLs, CENO-1B-1M scored highest within 1 kb of the transcription start site, but that advantage did not extend to the most distal stratum, where HyenaDNA-Medium-450k and Evo 2 7B led among the ten displayed models (Figure 3f). Structural-variant deletion scoring was more uniformly favorable to CENO, with all three CENO-1B stages occupying the leading AVG2 positions (Figure 3g). By contrast, metabolic-pathway performance remained distributed across the CENO and GENERator families (Figure 3h).

The supplementary analyses separated broad transfer from subgroup-specific leadership (Supplementary Fig. S5). CENO-1B had the strongest family-level ClinVar score, followed by Evo 2 7B and CENO-600M (Supplementary Fig. S5a). OmniReg-GPT led each displayed clinical-group F1 comparison, while the CENO-1B checkpoints showed marked group dependence (Supplementary Fig. S5b). The family-level advantage therefore reflected breadth across ClinVar tasks, not uniform superiority within every clinical category.

PromoterAI reinforced this distinction. AlphaGenome RNA-seq led the aggregate summary, followed by Evo 2 40B. The three CENO-1B stages formed the next group, with mean AUROC values confined to 0.563–0.568 (Supplementary Fig. S5c). The same top-two separation persisted for UKBB proteome under-expression, where the CENO-1B stages clustered behind AlphaGenome RNA-seq and Evo 2 40B (Supplementary Fig. S5d). These results support transfer across expression-perturbation tasks, while showing that model ordering remained task-specific.

On BRCA and enhancer-gene prediction tasks, Evo 2 models led, and CENO-1B-1M remained fourth for BRCA2 noncoding variants. All three CENO-1B stages also remained within the top eight for BRCA1 coding variants (Supplementary Fig. S5e,f). In enhancer-gene prediction, CENO-600M-1M ranked third behind Evo 2 40B and Genos-10B-v2 (Supplementary Fig. S5g). Performance therefore depended on more than model scale or nominal context length. TraitGym made this context dependence explicit (Supplementary Fig. S5h). CENO-1B-131k remained stable from 8k to 131k input, whereas CENO- 1B-1M peaked at 32k and declined at 131k. The benefit of longer input therefore depended on both the checkpoint and benchmark.

Together, these analyses show that CENO remained competitive across diverse task types. It ranked near the leading models across heterogeneous local and long-window effects, but its relative advantage varied with assay, variant class and context.

### 2.4. CENO generation recovers gene-scale sequence and supports structure-guided design

A generative genome model should do more than score existing sequence: it should propose sequence that satisfies a stated objective. Having established that CENO carries long-range signal in its representations, we tested its generative distribution in two settings that separate reconstruction from design. Withheld partial-gene continuation asked whether the model reproduces natural sequence it has not seen. Structure-guided insertion design then asked whether the model can supply long candidate sequences whose predicted chromatin effect survives evaluation by a model that took no part in selecting them.

We first asked whether CENO reproduces natural sequence that was withheld from it. Genes were drawn from eight species spanning eukaryotic, bacterial and archaeal domains; the first 30%, 50% or 80% of each gene was supplied as a prompt and the model generated the remainder, which was compared with the natural sequence (Methods 4.13). Figure 4b reports the 80% prompt, the most constrained setting. Averaged over the eight species, CENO-1B reached 44.1% recovery against 43.7% for Evo 2 7B, a model with roughly seven times the parameters, and exceeded Evo 2 1B by 6.9 to 7.6 percentage points in every phylum. CENO-1B gave the highest recovery of the eight models in all three eukaryotic species and in *B. subtilis* and *S. solfataricus*, exceeding Evo 2 7B by 4.6 points on eukaryotic and 2.7 points on archaeal genes. On bacterial genes it trailed Evo 2 7B by 5.4 points, with Evo 2 7B leading in *E. coli* and *S. coelicolor*. Recovery rose with model scale at every prompt length (CENO-300M 34.1%, CENO-600M 40.6%, CENO-1B 44.1% at the 80% prompt). Exact recovery is not the only meaningful outcome, because a model may generate sequence that differs from the natural allele while retaining related function.

**Figure 4.**
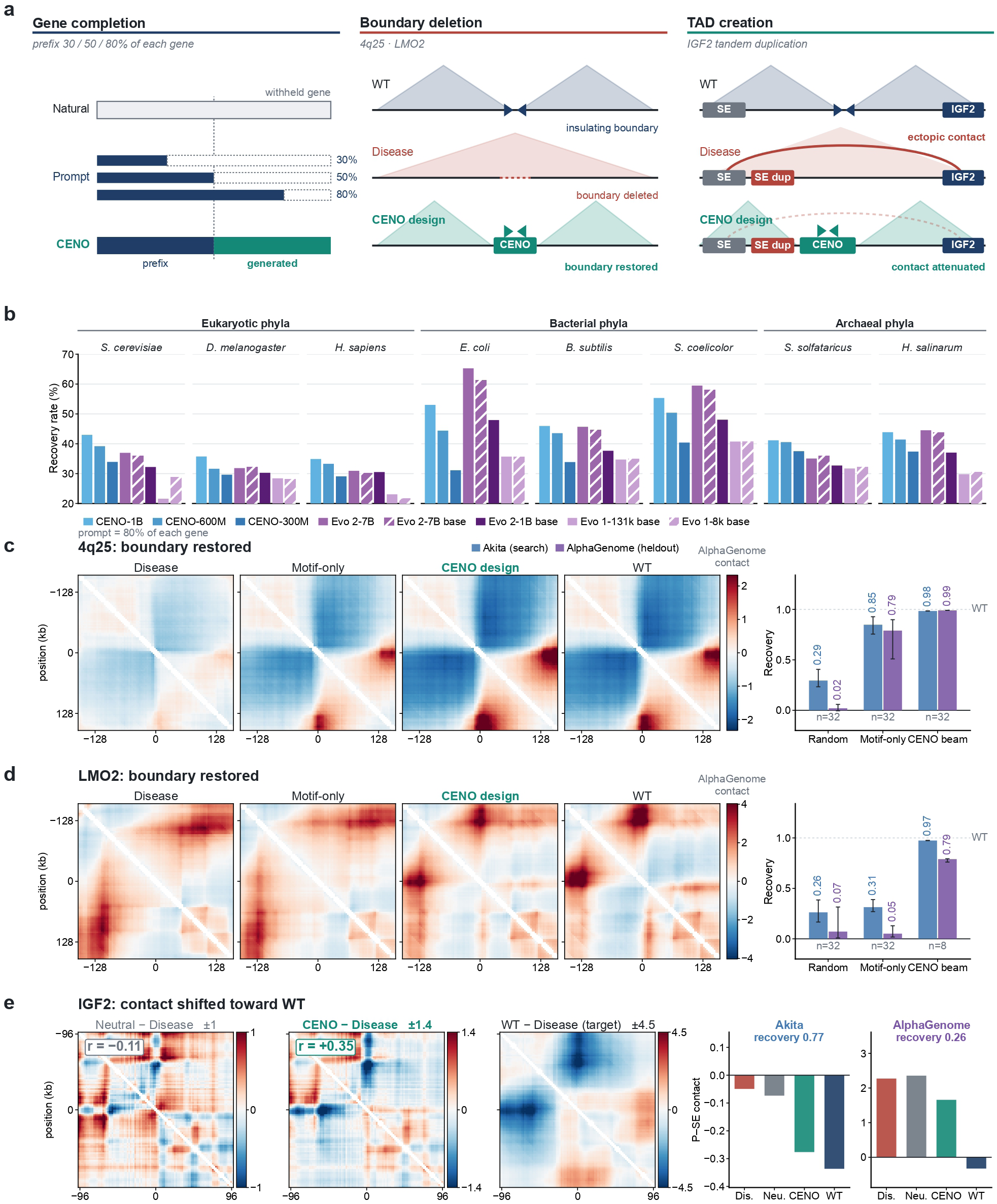
Gene-scale sequence recovery and structure-guided insertion design. **a,** The three tasks. *Gene completion*: the first 30%, 50% or 80% of a withheld gene is supplied as a prefix and the remainder generated, then compared with the natural sequence. *Boundary deletion*: a deleted insulating boundary is rebuilt by a generated insertion. *TAD creation*: a tandem duplication places a duplicated super-enhancer (SE) in the same domain as IGF2, creating an ectopic promoter–super-enhancer contact that a generated insertion is intended to re-insulate. **b,** Gene recovery rate at the 80% prompt across eight species in three domains for eight models, in fixed legend order; hatched bars are base checkpoints. The axis is truncated at 20%, so bar length is not proportional to value, and two bars fall below that floor and are drawn as clipped stubs. The 30% and 50% prompt settings are in the Supplementary material. **c,d,** 4q25 and LMO2. Held-out AlphaGenome contact maps for the disease allele, a representative motif-only insertion, a representative CENO insertion and the wild type, shown on one color scale per locus with the near-diagonal band masked. Right, median recovery per candidate group under the search model (Akita [32]) and the held-out model (AlphaGenome); whiskers span the interquartile range, the dashed line marks wild type, and **d** uses the key in **c**. Beam candidates share a search lineage, so their narrow interval is not independent replication. **e,** IGF2 difference maps relative to the disease allele, each scaled to its own limit (printed after the title); *r* is the correlation with the wild-type change. Right, raw promoter–super-enhancer (P–SE) contact per condition under each model with the recovery it implies; the two axes differ roughly tenfold and are not comparable. Throughout, *n* denotes the number of candidates. Loci, insertion lengths, cell types and recovery definitions are in Methods 4.14.

We next asked whether the pretrained model could supply long insertions with a specified effect on predicted chromatin contacts. Two loci provided a boundary-deletion setting in which the disease mechanism is loss of insulation: deletion of a TAD boundary at 4q25 causes PITX2-related cardiac defects [33], and deletion of a boundary adjacent to LMO2 activates the proto-oncogene [34]. At each locus Akita ranked CENO-generated, motif-only and matched-random insertions under oracle-guided beam search; the candidate set was then frozen and scored with AlphaGenome across five positional shifts, with AlphaGenome taking no part in selection (Methods 4.14). Recovery was scaled so that the wild-type map is 1 and the deletion map is 0.

At 4q25, complete 15,312-bp CENO insertions reached median recovery of 0.984 under Akita and 0.990 under held-out AlphaGenome (*n* = 32; Figure 4c). The search used an H1-hESC Akita target and the held-out evaluation an H9 ventricular-cardiomyocyte AlphaGenome track, so this transfer crosses both model and cell type. Motif-only insertions also transferred, reaching 0.848 and 0.790, whereas matched-random insertions reached 0.295 under Akita and 0.020 under AlphaGenome (*n* = 32 each). Complete generated insertions therefore retained a strong boundary-associated effect at this locus, where local motif grammar was already sufficient.

At LMO2, where both models score the same GM12878 target, 26,629-bp CENO insertions reached median recovery of 0.974 under Akita and 0.783 under AlphaGenome (*n* = 8; Figure 4d). Motif-only insertions reached 0.314 and 0.050 and matched-random insertions 0.262 and 0.070 (*n* = 32 each), so neither control reproduced the cross-model effect. Two features of the candidate set bound this result. The eight beam candidates share a search lineage and are 92% identical to one another on average, against 81% at 4q25 and 25–28% within the control groups, so they are not eight independent solutions. CENO sequences sampled without oracle ranking reached median held-out recovery of −0.016 at LMO2 and −0.019 at 4q25, indistinguishable from matched-random insertions. The boundary-restoring effect therefore requires the generative model together with oracle-guided search, rather than the pretrained sampling distribution alone.

A tandem duplication at IGF2 provided a harder setting in which the mechanism is gain of an ectopic contact rather than loss of insulation: duplication brings a super-enhancer into the IGF2 domain and activates the gene by enhancer hijacking [35]. Because the rearrangement creates a neo-domain, we scored the promoter–super-enhancer (P–SE) contact directly, with one definition applied to both models and referenced to a length-matched neutral insertion (Methods 4.14). A selected 15,312-bp CENO insertion recovered 0.26 of the neutral-to-wild-type gap under held-out AlphaGenome and 0.77 under Akita (Figure 4e). The two values differ because the models disagree about the size of the lesion, not about the effect of the insertion: AlphaGenome predicts a strongly positive ectopic P–SE contact in the disease allele (+2.27 against −0.33 for wild type) whereas under Akita the same contact remains depleted (−0.05 against −0.34), leaving a denominator 10.2 times smaller, and in absolute terms the insertion moves the contact 3.4 times further under AlphaGenome. The change was spatially concordant in both models: the difference map relative to the disease allele correlated with the wild-type-versus-disease difference at *r* = +0.35 under AlphaGenome and *r* = +0.42 under Akita, whereas the neutral insertion showed no such correspondence (*r* = −0.11 and +0.11). Across 38 candidates from five sampling strategies that completed cross-model evaluation, the highest held-out P–SE recovery was 0.26, so this value reflects a limit of the present search rather than the choice of one candidate. The generated insertion therefore shifted the ectopic contact toward wild type in the correct positions and at roughly a quarter of the required amplitude, without restoring the wild-type map.

These experiments separate two uses of the CENO generative distribution. CENO recovered natural gene continuations at a level competitive with a model seven times its size and clearly better at matched parameter count, and, coupled to an oracle-guided search, supplied long insertions whose predicted boundary-associated effects survived evaluation by a second structure model and, at 4q25, a second cell type. In the duplication setting the selected candidate moved the target contact in the correct direction but only partially, and the two structure models disagreed substantially about the magnitude of the target.

### 2.5. MSA-context post-training improves variant scoring across evolutionary scales

To adapt CENO to evolutionary context, we post-trained the pretrained long-context DNA backbones on packed examples built from real multiple sequence alignments (MSAs); we refer to the resulting MSA-adapted models as CENO-P. Each example contains a target sequence together with homologous rows. The post-training stack interleaves row-local layers, which process each sequence independently, with fusion layers that allow homologous rows to condition the target representation (Figure 5a; Methods 4.9). Variant-effect prediction then uses the same zero-shot interface as pretraining: reference and mutant alleles are scored under the same MSA context, and the likelihood delta is used as the variant score (Figure 5b; Methods 4.11). The post-training objective is a causal next-token loss over packed real MSA contexts (Methods 4.10).

**Figure 5.**
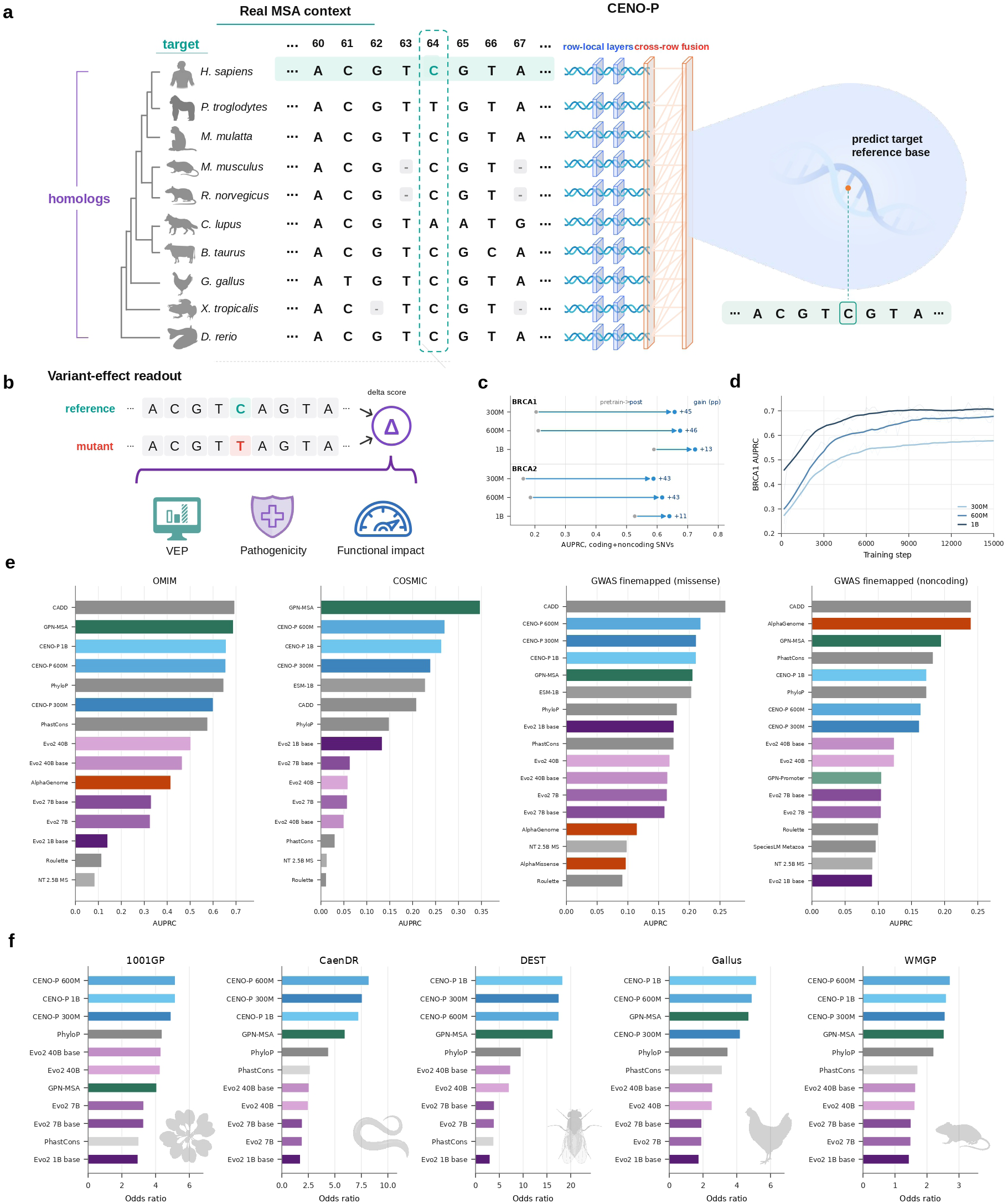
CENO post-training with real MSA context. **a,** CENO packs a target sequence with homologous rows and explicit row identities. **b,** Variants are scored by the reference-versus-mutant likelihood delta under the same MSA context. **c,** Matched CENO pretraining and post-training checkpoints on BRCA1 and BRCA2. **d,** BRCA1 zero-shot AUPRC over post-training. **e,** Disease and GWAS fine-mapping benchmarks with all plotted models; post-trained checkpoints are labeled CENO-P. **f,** Cross-species Fisher odds-ratio enrichment benchmarks.

We first asked whether post-training improved each backbone relative to its own pretrained checkpoint. This matched comparison isolates the effect of MSA-context adaptation from differences between model families. On BRCA variant-effect benchmarks following the GPN-MSA evaluation protocol [13], post-training increased AUPRC by 45.3%, 46.4% and 13.5% for the 300M, 600M and 1B models, respectively (Figure 5c). The corresponding BRCA2 gains were 42.6%, 43.1% and 11.3%. Thus all three CENO backbones gained variant-effect signal in their CENO-P form after exposure to real homologous context.

These gains accrued over the course of post-training rather than appearing only at the final checkpoint. Tracking BRCA1 zero-shot AUPRC over the first 20k steps with the likelihood-delta scoring interface (Methods 4.11), we found that performance climbed steeply during the first few thousand steps and then plateaued at all three scales. Thus, most of the variant-effect signal was acquired early in post-training (Figure 5d).

We then evaluated the post-trained checkpoints on the human disease and fine-mapping tasks used in the GPN-Star benchmark panel [14], together with the plotted baselines (Figure 5e). These tasks comprise COSMIC somatic missense variants [36], OMIM regulatory variants using the TraitGym-processed Mendelian dataset [37, 23], and coding and noncoding GWAS fine-mapped variants from UK Biobank fine-mapping resources [38, 39, 23]. Among the models retained in this panel, the best CENO-P ranked third on OMIM (AUPRC 0.657), second on COSMIC (0.270), second on GWAS fine-mapped missense variants (0.219), and fifth on GWAS fine-mapped noncoding variants (0.173). Across these human-disease tasks, CENO-P was competitive with strong specialized and conservation-based genome-wide scorers.

CENO-P’s clearest advantage was in cross-species conservation. Following the GPN-Star cross-species benchmark construction [40, 41, 42, 43, 44], we scored Fisher odds-ratio enrichment on five species panels (Figure 5f; Methods 4.12). A CENO-P checkpoint ranked first among the plotted models on all five panels (1001GP, CaenDR, *D. melanogaster*, Gallus and WMGP), with odds ratios of 5.16, 8.21, 18.30, 5.16 and 2.72, respectively, surpassing genome language models an order of magnitude larger as well as alignment-based conservation scores. This uniform lead across evolutionarily distant clades showed that MSA-context post-training transferred broadly across the tree of life and was most effective when homologous rows were available during both adaptation and variant scoring.

### 2.6. CENO enables mouse cortical cell-type-specific enhancer prediction and design

We investigated cell-type-specific enhancer prediction and design with CENO using a mouse motor cortex scATAC-seq dataset. The workflow couples a CENO-based accessibility oracle with a conditional sequence generator (Figure 6a). Candidate CREs were first derived from mouse cortex scATAC-seq peaks and organized into a cell-type-by-accessibility matrix. The oracle was trained to predict subclass-resolved ATAC activity from the same input DNA sequence, and the resulting sequence-to-function model was used as a reward model for conditional enhancer generation. The generator was first adapted by supervised fine-tuning (SFT) to learn cell-type-conditioned CRE sequence distributions and was then further optimized by reinforcement learning (RL) to increase predicted target-cell activity while penalizing off-target activity.

**Figure 6.**
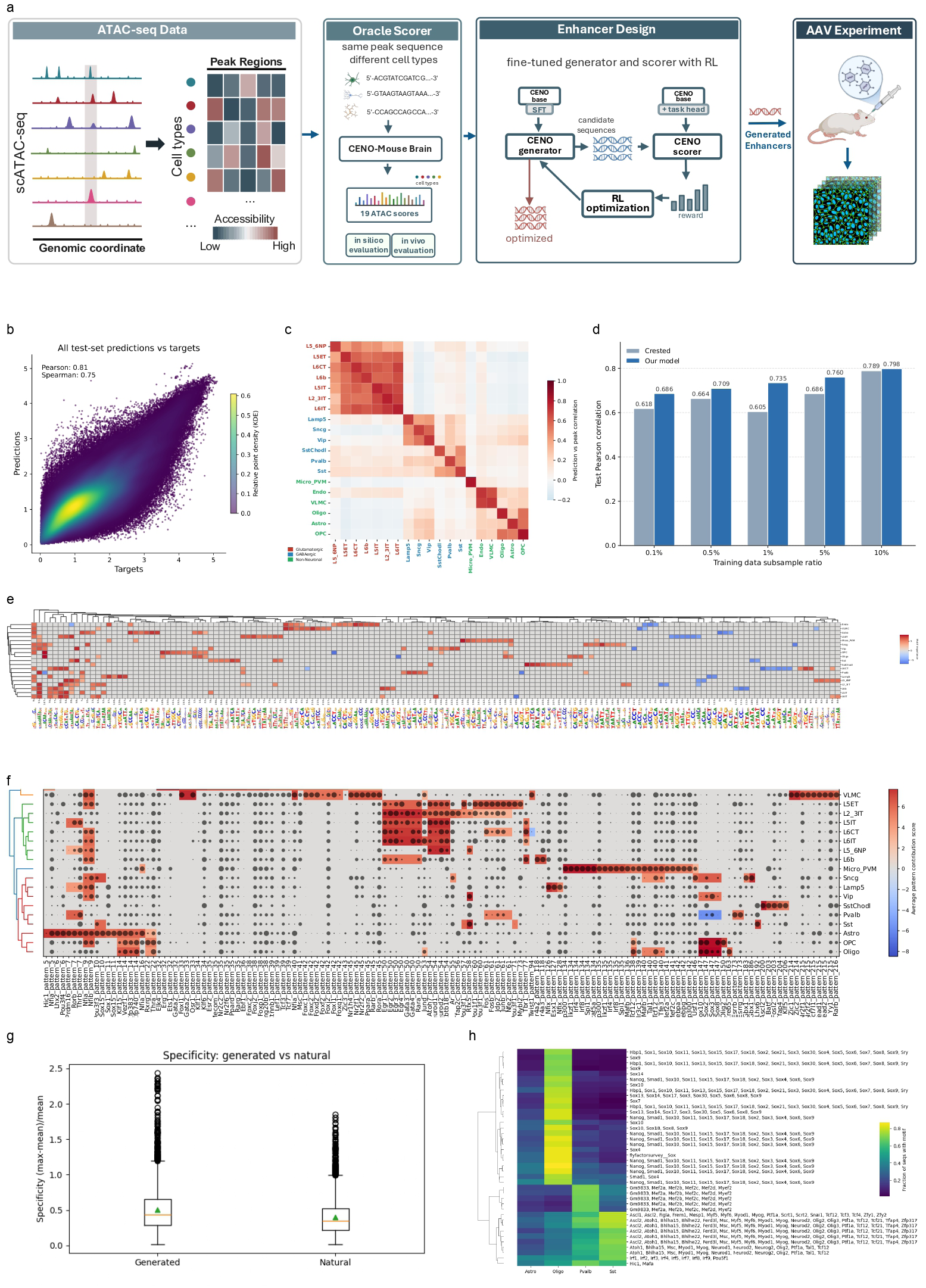
Cell-type-specific enhancer prediction and design. **a,** Overview of oracle-guided cell-type-specific CRE generation. **b,** Density scatter plot of oracle predictions and observed chromatin accessibility on held-out test peaks. **c,** Cross-cell-type correlation matrix between predicted and observed chromatin accessibility. **d,** Data-efficiency comparison between the CENO-based oracle and the CREsted baseline [45] across training-set subsampling fractions, evaluated by test Pearson correlation. **e,** TF-MoDISco motif atlas derived from oracle contribution scores. **f,** Joint visualization of motif contribution scores and matched TF RNA-seq expression. **g,** Predicted specificity of generated and natural CREs. **h,** Motif enrichment across Astro, Oligo, Pvalb and Sst generated sequences.

We first trained a CENO-based peak regression oracle on 440,993 consensus peaks and then fine-tuned it on 73,323 cell-type-specific regions. To quantify oracle performance, we evaluated predictions on chromosome-held-out consensus and cell-type-specific test peaks (Figure 6b; Supplementary Fig. S7b). For each test region, the oracle produced accessibility scores across mouse cortex subclasses. We flattened predictions and targets across peak–cell-type pairs, excluded entries with zero observed accessibility, and compared log1p-transformed predictions with log1p-transformed peak heights. The oracle achieved a Pearson correlation of 0.81 and a Spearman correlation of 0.75 on these nonzero pairs, with the density scatter plot showing a clear monotonic relationship between predicted and observed accessibility.

We further asked whether oracle prediction channels preserved cell-type identity. For each predicted cell type and each observed cell type, we computed Pearson correlation across cell-type-specific test peaks (Figure 6c). The strongest correlations were concentrated along the diagonal, indicating that each prediction channel best matched its corresponding target cell type. Block-wise correlations within glutamatergic, GABAergic and non-neuronal groups further reflected shared accessibility programs among related cell types.

We then examined whether the oracle learned interpretable regulatory grammar. Nucleotide-level contribution scores were computed by in silico mutagenesis (ISM), which measures the change in oracle prediction after single-nucleotide substitutions. Recurrent high-contribution sequence patterns were then identified and summarized as a cell-type-by-motif importance matrix for the 2,000 most cell-type-specific regions (Figure 6e). The resulting motif atlas showed structured, cell-type-associated motif usage across cortical subclasses, suggesting that oracle predictions are supported by recognizable sequence features rather than only generic accessibility-associated signals.

To connect these sequence features with cell-type biology, we jointly compared motif contribution scores with matched TF expression from RNA-seq (Figure 6f). Several high-contribution motifs showed concordant cell-type-specific TF expression patterns. For example, NFI-family motifs were prominent in astrocyte-related non-neuronal classes, whereas SOX-family motifs were enriched in Oligo/OPC-associated programs. These results indicate that the oracle captures sequence features aligned with cell-type-specific TF programs.

Having established the oracle, we next evaluated the sequences produced by oracle-guided generation. Generated sequences were sampled from the final GRPO generator, which was optimized from the SFT model using the tail-only oracle MinGap reward without demonstration injection (Methods 4.15). RL-generated sequences showed higher oracle-predicted target-cell scores than natural CREs for Astro, Oligo and Sst, but not for Pvalb (Figure S8). We then quantified predicted cell-type specificity for generated and natural CREs (Figure 6g). Specificity was defined as the maximum predicted activity across cell types divided by the mean predicted activity across cell types, measuring whether activity is concentrated in one cell type rather than broadly distributed. Generated sequences showed an upward shift in predicted specificity relative to natural CREs, indicating that oracle-guided generation enriched for more cell-type-selective candidate enhancers.

Finally, we assessed whether generated sequences retained recognizable cell-type-associated motif grammar (Figure 6h). Motif enrichment analysis was performed on sequences generated for Astro, Oligo, Pvalb and Sst targets, and the fraction of sequences containing each motif family was summarized across target classes. Generated sequences showed target-dependent motif composition, including SOX-family enrichment in Oligo-targeted designs and MEF2-family enrichment in Pvalb-targeted designs. Thus, conditional generation increased oracle-predicted specificity while preserving interpretable motif features associated with the target cell type.

Overall, this workflow provides an oracle-guided route from mouse cortex accessibility modeling to enhancer-code interpretation and cell-type-specific CRE generation. It demonstrates CENO’s utility for sequence-to-function prediction and regulatory sequence design.

## 3. Discussion

CENO was developed from the view that genome modeling should become a world-modeling problem. DNA is not only a string to be classified, but a generative substrate whose local syntax, distal regulatory structure, evolutionary history and designable function are coupled across scales. In this operational sense, a genomic world model should support multiple modes of use within one backbone: reading existing sequence, scoring counterfactual variants, reconstructing or generating plausible sequence continuations, conditioning on evolutionary context and optimizing new sequences under functional constraints. CENO connects these modes through a single autoregressive backbone, long-context continuation, evolutionary MSA post-training and oracle-guided regulatory design.

The term world model should be interpreted operationally rather than metaphysically. CENO does not claim to simulate complete cellular state, organismal phenotype or physical 3D genome dynamics. Instead, it models the sequence-native world of genomic constraints: local nucleotide grammar, gene-scale organization, distal regulatory context, homologous evolutionary evidence and designable regulatory sequence space. This framing is useful because it turns a broad ambition into testable capabilities. A genomic world model should be judged by whether it can retrieve distal information, form long-context representations, score sequence interventions, recover withheld sequence, use homologous context and generate sequences under functional objectives.

The architecture and curriculum of CENO were designed to separate model capacity, context length and data composition. The hybrid Mamba–attention–MoE backbone combines efficient long sequence mixing, explicit content-based interactions and sparse expert capacity. The staged curriculum first learns broad genomic sequence grammar with an 8k-token context, then shifts toward eukaryotic, transcript, splice and regulatory sequence sources, and finally extends context to 131k and 1M tokens with complete eukaryotic-genome windows. This separation is important because short-range and long-range genomic capabilities are not identical. Local motif grammar and coding syntax can often be captured from short windows, whereas distal retrieval, chromatin-boundary-associated representations and gene-scale continuation require both longer context and sufficient model capacity. In the world-model framing, increasing context length is not itself the goal; the goal is to make longer sequence context usable by the model state.

The long-context analyses indicate that CENO does more than accept longer inputs. In the DNA needle assay, mutating a distant inserted sequence changed terminal base predictions across 32k, 64k and 131k contexts, showing that distal sequence can influence the local probability distribution. In the TAD analyses, 131k continuation strengthened within-boundary relative to across-boundary attention in the 600M and 1B models and improved frozen-state boundary probes across human and mouse settings. These results suggest that long-context continuation changes the internal representation of genomic sequence, rather than merely increasing the configured context length. At the same time, these assays should be interpreted as diagnostic readouts. TAD-scale attention and linear probes provide evidence for chromatin-boundary-associated sequence representations, but they do not imply that the model reconstructs a complete physical contact map or replaces supervised 3D genome prediction.

The cross-setting behavior is an important part of this interpretation. The TAD attention and frozen-probe signals are not restricted to one displayed locus or one cell type: they are observed across multiple human cell types and mouse cell or tissue settings. This suggests that long-context continuation can produce reusable sequence-level boundary representations rather than a single-context artifact. However, the effect is scale dependent. The 600M and 1B models show the clearest gains, whereas the 300M model is weaker in several chromatin-boundary readouts. This pattern suggests that long-context genomic reasoning may require a capacity threshold: increasing context length alone is not sufficient unless the model has enough representational capacity to use the additional sequence.

The broader zero-shot benchmarks reinforce the need for multi-axis evaluation. Variant-effect prediction, molecular fitness, regulatory-impact tasks, structural-variant readouts and sequence generation probe different aspects of genomic modeling. CENO is competitive across many of these settings and strong in several long-context and evolutionary-context analyses, but no single benchmark family is sufficient to define general genomic ability. Human disease and fine-mapping benchmarks remain especially challenging, and task-specific or phylogeny-informed methods can remain stronger in some settings. This fragmentation should not be viewed only as a limitation of CENO. It reflects the current state of genomic foundation-model evaluation, where short-range regulatory grammar, protein or RNA fitness, noncoding variant effects, chromatin-scale structure and generation fidelity are often measured by different tasks with different inductive biases.

The world-model framing changes how genomic foundation models should be benchmarked. A model that performs well on local variant-effect prediction may still fail to use distal context. A model that accepts long inputs may still fail to generate coherent gene-scale continuations. A model that generates plausible DNA may still fail under functional design constraints. We therefore view genomic world-model evaluation as a unified benchmark paradigm rather than a single task family. Retrieval assays test whether distant sequence is accessible to the model state. Chromatin-boundary attention and frozen probes test whether long-context representations carry locus-scale structure. Variant-effect benchmarks test counterfactual perturbation sensitivity. Partial-gene continuation tests whether the learned distribution can reconstruct species-appropriate sequence organization. MSA-conditioned scoring tests evolutionary reasoning. Oracle-guided enhancer generation tests whether the same backbone can be adapted from interpretation to design.

Long-sequence generation provides a useful counterpart to variant-effect prediction. VEP asks whether a likelihood-based model assigns meaningful score changes to local perturbations of an observed reference sequence. Generation benchmarks ask a different question: given a partial gene or genomic prompt, can the model recover the withheld continuation with high similarity and species-appropriate sequence structure? This is a reconstruction-style test of gene-scale sequence understanding. It requires the model to maintain local composition, coding and noncoding grammar, species-specific sequence statistics and longer-range organization during autoregressive decoding. The improvement of generation recovery with larger CENO models and later whole-genome long-context continuation therefore supports the view that the later curriculum strengthens gene-level sequence modeling, not only inference throughput or variant scoring. This benchmark should nevertheless be interpreted as reference recovery rather than direct functional validation: recovering a natural continuation is evidence of sequence-distribution understanding, but it does not by itself prove that newly generated sequences have the intended biological function.

Evolutionary post-training adds a complementary source of information to single-sequence pretraining. By packing real MSA rows with explicit segment identities, CENO can condition the target sequence on homologous context while retaining a likelihood-delta interface for variant-effect scoring. The matched pretraining-versus-post-training comparisons on BRCA1 and BRCA2 show that the same backbone gains substantial variant-effect signal after exposure to real homologous context. The strongest pattern appears in cross-species odds-ratio enrichment benchmarks, where CENO benefits from seeing homologous rows during both adaptation and scoring. This suggests that explicit evolutionary context provides information that is not fully captured by isolated-sequence likelihoods or by conservation labels alone. At the same time, the benefit depends on the availability, quality and relevance of homologous alignments. Sparse clades, poorly aligned regions and species without dense alignment resources remain important future test cases.

CENO also connects interpretation to sequence design. In the mouse cortex CRE workflow, a CENO-based accessibility oracle provides a cell-type-specific sequence-to-function interface, supervised fine-tuning teaches the generative model to condition on raw text cell labels, and GRPO uses oracle feedback to shift generated sequences toward target-cell-type activity. The resulting sequences show substantial oracle-score increases for three of the four target classes and retain recognizable target-associated regulatory motifs, including NFI-family motifs for astrocyte targets, SOX-family motifs for oligodendrocyte targets and MEF2-family motifs for Pvalb-associated designs. This supports the use of CENO as a design backbone rather than only a sequence scorer. Importantly, the design setting is different from the long-sequence generation benchmark. Generation recovery tests whether the pretrained model can reconstruct natural sequence continuations, whereas enhancer design tests whether the model can be guided toward new sequences satisfying a functional objective.

Oracle-guided regulatory design also introduces limitations that should be made explicit. Reinforcement learning can exploit weaknesses in the oracle, especially when generated sequences move away from the natural CRE distribution. Motif enrichment, target-associated TF programs and retrospective oracle validation provide useful evidence that the optimized sequences remain biologically plausible, but they are not substitutes for prospective experimental validation. Designed enhancers should therefore be treated as prioritized candidates for MPRA, AAV or other wet-lab assays, not as validated functional elements. Future closed-loop design should incorporate experimental feedback, distribution-shift checks, motif-level and syntax-level constraints, and negative controls designed specifically to detect oracle hacking.

Several broader limitations remain. First, the current long-context diagnostics focus on retrieval, chromatin-boundary-associated attention and TAD-boundary linear probes. These do not cover all forms of long-range genome regulation. Enhancer–promoter communication, locus control regions, insulation, structural variation and context-dependent transcriptional regulation will require additional assays and task-specific readouts. Second, downstream performance is not strictly monotonic with model size. Larger models generally benefit long-context and generation recovery, but smaller models can gain substantially from evolutionary post-training in some variant benchmarks. This suggests that data curriculum, context length, post-training objective and architecture may matter as much as total parameter count. Third, MSA post-training currently depends on real homologous rows; generated MSA contexts, low-resource species and alignment-free evolutionary conditioning remain open directions. Fourth, sequence generation benchmarks based on reference recovery measure distributional reconstruction rather than function. They are valuable for testing gene-scale sequence understanding, but should be combined with functional or experimental assays when making biological claims. Fifth, the current design workflow uses an oracle learned from available mouse-cortex accessibility data; extending genomic world models toward broader phenotype spaces will require richer oracles, perturbational measurements and prospective experimental feedback.

These limitations point to a broader path forward. Long-context genomic world models can serve as a common substrate for three increasingly functional uses: single-sequence genome modeling, evolutionary sequence interpretation and programmable sequence design. In this view, pretraining learns a short-to-long representation of genome sequence, MSA post-training adds evolutionary constraint, and oracle- or experiment-guided optimization turns the model into a design engine. Future CENO extensions could incorporate richer alignment generation, tissue- and perturbation-specific oracles, prospective validation of designed regulatory elements, and closed-loop variant-to-design workflows. More generally, our results suggest that genomic foundation models should be developed as world models of genome sequence space: models that retain nucleotide-resolution grammar, route information across regulatory and chromatin-scale distances, recover gene-scale sequence organization across species, condition on evolutionary evidence and adapt these representations to functional design.

## 4. Methods

### 4.1. Model architecture

CENO is a family of causal autoregressive genomic language models with a hybrid Mamba-2, attention and mixture-of-experts backbone [46, 47]. We trained three scales with approximately 300M, 600M and 1B total parameters, corresponding to approximately 100M, 200M and 400M active parameters per token. All models use a byte-level DNA tokenizer with a 512-entry vocabulary. Input and output embeddings are untied, no learned absolute position embeddings are used, and the maximum configured context length is 1,048,576 tokens.

The backbone contains Mamba-2, attention and MoE layers. Mamba-2 layers use state dimension 128 and head dimension 64. Attention layers provide explicit content-based interactions, and MoE layers use eight experts with router top-*k* = 2. The 300M, 600M and 1B models use 9, 20 and 38 layers, respectively. The 600M configuration replaces the first MoE position with a dense MLP block for training stability.

### 4.2. Pretraining data and schedules

CENO was pretrained on OpenGenome2, the same broad genomic corpus used by Evo 2 [7]. The tokenized corpus spans GTDB/IMG prokaryotic genomes, metagenomes, viral genomes, organelle genomes, eukaryotic genic windows, mRNA, splice/promoter windows, ncRNA, promoter windows and complete eukaryotic genomes. We retained the Evo 2-style train, validation and test split supplied with the tokenized corpus. Evo 2/OpenGenome2 safety exclusions, including the exclusion of eukaryotic-host viruses, were retained during preprocessing.

Pretraining used causal next-token prediction with a staged context-length curriculum. Stage I trained at 8,192 tokens with a broad OpenGenome2 mixture. Stage II kept the 8,192-token context and increased the eukaryotic, transcript, splice and regulatory components. The 300M, 600M and 1B models used Stage I/II budgets of 1.0T/0.5T, 2.0T/1.3T and 3.5T/2.5T tokens, respectively. Stage III continued from the Stage II checkpoint at 131,072 tokens for 500B tokens. Stage IV continued from the Stage III checkpoint at 1,048,576 tokens for an additional 500B tokens. Model-specific sample counts and batch settings are listed in Supplementary Table S1.

Stage I and Stage II mixtures were selected with constrained RegMix [48]. For Stage I, 512 candidate proxy models spanning seven source categories were generated under a broad eukaryotic/prokaryotic balance constraint and evaluated on a selected subset of the VEP benchmarks. A LightGBM response model [49] was trained to map mixture weights to mean benchmark performance, scored 200,000 simulated mixtures, and produced the final Stage I weights by averaging the top 128 predicted mixtures. Stage II used the same selection procedure, using 507 candidate proxy models over nine source categories, 100,000 simulated mixtures and stronger eukaryotic, transcript, splice and regulatory weighting constraints. The fixed Stage I/II production weights are summarized in Supplementary Table S2.

Stage III and Stage IV used a long-context continuation mixture designed to preserve the Stage II eukaryotic-dominant balance while allocating part of the eukaryotic component to complete eukaryotic genomes. It retained short-window, transcript, splice, regulatory and microbial sources and added Animalia, Plantae, Fungi, Protista and Chromista complete-genome sources sampled into 131k and 1M windows. At the grouped level, Stage III/IV assigned 32.32% to prokaryotic, metagenomic, viral and organelle sources and 67.68% to eukaryotic, transcript, regulatory and complete-genome sources.

Optimization used Adam with *β*_1_ = 0.9, *β*_2_ = 0.95, weight decay 0.1, gradient clipping at 1.0, cosine learning-rate schedules and FP8 mixed precision. Attention and hidden dropout were set to zero. Repeat-masked lowercase bases were retained in the input sequence and assigned loss weight 0.1 in the training objective. Long-context stages used context parallelism to shard the sequence dimension. Training loss, validation loss and pretraining throughput summaries were computed directly from Megatron logs for the reported stages.

### 4.3. Inference throughput

Inference throughput was evaluated with random DNA prompts on a single GPU. Two regimes were measured. In input-length scaling, output length was fixed at 512 tokens and prompt length was varied up to 131,072 tokens where the model supported the requested context. In generation-length scaling, prompt length was fixed at 1,024 tokens and generated length was varied up to a total sequence length of 131,072 tokens. Throughput was reported as generated tokens per second per GPU for each condition. Out-of-memory or runtime failures at a requested input length were recorded as failed runs and retained in the throughput grid.

CENO checkpoints were evaluated using a modified vLLM backend with support for the CENO architecture. Evo-family baselines were evaluated with their supported inference backends and model-specific code paths on the same hardware. For the Figure 1f input-scaling display, raw benchmark summaries were retained for every condition.

### 4.4. Needle-in-a-haystack assay

We used a needle-in-a-haystack assay adapted from the Evo 2 study [7] to test whether sequence information placed far from the prediction site affected terminal predictions. For each sample, a random 100-bp DNA needle was inserted into a random DNA haystack at a specified depth, and an identical 100-bp query was appended to the end of the prompt. Main Figure 2 reports test contexts of 32k, 64k, 131k and 1M bp at insertion depths of 10%, 50% and 90%. We evaluated three independently generated samples for every model–context–depth condition.

Retrieval was quantified using diagonal categorical-Jacobian-style perturbations. At each needle position, we first recorded the A/T/C/G log-probability vector at the aligned query position. We then substituted the inserted base with each of the other three nucleotides and recomputed the corresponding query-position vector. The position score was the mean Euclidean distance between the original and perturbed four-base vectors over the three substitutions, and the retrieval score was the mean of the 100 position scores.

In positive samples, the inserted needle was identical to the terminal query. For shuffled-null samples, the inserted needle was randomly permuted while the terminal query was held fixed. Null controls were evaluated at 10% and 100% insertion depths, with three independently generated samples per condition. Matched unshuffled records were also retained to validate each null construction but were not included in the reported null statistics. A positive trial was counted as successful when its retrieval score exceeded the prespecified threshold of 0.8. Each displayed heatmap value is the mean of three positive trials. For each model-stage and test-context setting, the supplementary summary reports nine positive records (three depths by three trials) and six shuffled-null records (two depths by three trials).

### 4.5. Zero-shot TAD attention analysis

We used TAD-boundary loci as a zero-shot readout of long-context routing. No boundary labels were used to train or adapt the models for this analysis. Source boundary files were obtained from the 4DN Project. Boundary intervals labeled Strong with score at least 0.5 were retained. We centered 131,072-bp windows on retained boundaries, sampled up to 128 positive windows per source and generated same-length negative windows at least 50,000 bp from any retained boundary window using seed 42. Negative rows were sampled from genomic windows that did not overlap the retained annotations. The main Figure 2b,c scores were computed from boundary-centered positive windows.

For CENO, maps were computed at bin resolution rather than from full token-token attention tensors. At each selected attention layer, Q and K states were mean-pooled inside 128-bp bins, converted to causal QK softmax attention per head, averaged over heads and then averaged over selected layers. Each centered matrix was partitioned into disjoint segment masks. TAD-map panels display log_2_ enrichment relative to the across-boundary baseline. The primary score was the log_2_ ratio of mean within-boundary attention to mean across-boundary attention. Context-scaling panels compared matched Stage II 8k-token baselines and 131k-token continuation settings for the same locus.

Each extracted locus has complete controls for CENO 300M, 600M and 1B Stage II 8k; CENO 300M, 600M and 1B 131k; Evo 2 7B base 8k; and Evo 2 7B 131k. To summarize cross-context behavior, we computed species-level means and s.e.m. for each model family and context length. The summaries used the within/across-boundary score across five human cell types and five mouse cell or tissue settings. Human cell types were GM12878, H1-hESC, HFFc6, IMR-90 and K562. Mouse settings were brain, cerebellar granule neurons at postnatal day 11, double-positive thymocytes, Patski cells and splenic regulatory T cells.

### 4.6. Annotation-linked attention

Annotation-linked attention was evaluated at selected loci using CENO 1B with a 131k-token context. We examined cCRE, promoter, CTCF-bound, enhancer-like and CpG tracks. For each displayed case, one selected head or the median of a recorded selected-head set was aligned to genomic bins. Annotated bins were then compared with non-annotated background bins within the same local window. The track definitions were ENCODE cCRE GRCh38, UCSC CpG islands on hg38 and GENCODE v44 promoters within 2 kb of transcript starts. Public case labels, selected layer/head indices and bin counts are retained in the Figure 2d source tables. The attention profile was averaged across query positions, divided by the median local background attention and converted to log_2_ observed/background units. The positive profile was then compared with a matched negative profile for the same query and target-track relationship. Profiles were smoothed over five bins, detrended by a 39-bin-wide trend and robust-scaled against same-window background bins. One normalized unit denotes one background robust-scale unit above the background median after this matched-control and detrending procedure. Annotated bins are visible target track bins that exclude query bins; background bins are non-annotated, non-query bins in the same focus window. The selection audit required at least six annotated bins, at least one annotated-bin cluster and a centrality score of at least 0.45. Figure 2 reports a representative selection of these examples. The panel reports normalized attention profiles and bin-level summary bars.

### 4.7. TAD and CTCF linear probing

Frozen hidden states were extracted from each model and layer and used as fixed features for logistic-regression probes. The primary probe distinguished TAD boundary positions from matched negatives. A parallel CTCF probe distinguished CTCF motif positions from non-motif negatives and served as a local motif control. Probe positions were sampled from boundary-centered windows. Boundary positives were within 2,500 bp of the window center. Boundary negatives were at least 25,000 bp from the center and at least 200 bp from any CTCF motif center. CTCF positives used motif centers detected in each boundary-centered window, capped at 20 motifs per region. CTCF negatives were non-motif positions at least 200 bp from any motif center. Both samplers avoided the outer 64 bp of each window and used seed 2026.

Layer selection was fixed before cross-cell-type reporting. Human selected layers were chosen by H1-hESC boundary-probe performance. Mouse selected layers were chosen by brain boundary-probe performance. In-cell AUROC was the mean of five GroupKFold splits that left genomic-region groups out. Cross-cell-type transfer, where available, trained on all examples from one cell type and tested on another. Features were standardized within each training split. Logistic regression used *C* = 1.0, a maximum of 200 iterations and seed 2026. Human probe summaries used GM12878, H1-hESC, HFFc6, IMR-90 and K562. Mouse probe summaries used brain, cerebellar granule neurons at postnatal day 11, double-positive thymocytes, Patski cells and splenic regulatory T cells. Label-shuffle controls permuted labels 100 times at the selected boundary layer.

### 4.8. Matched-context variant-effect evaluation

To test whether long-context pretraining improved a downstream functional readout, we evaluated zero-shot variant-effect prediction on the TraitGym benchmark, which scores regulatory variants annotated as Mendelian or Complex traits. Variant effects were scored without task-specific training from the reference/alternate allele log-likelihood difference (Δ log *p*) under each model, and task performance was summarized as AUROC against the benchmark labels. Variants were stratified by variant–gene distance into <1 kb, 1–10 kb, 10–100 kb and >100 kb bins, and the Combined set pooled the Mendelian and Complex traits.

We defined a matched-context comparison in which model context and evaluation input length grew together: the 1B base (Stage II) checkpoint was scored with an 8k-token input window, and the 1B Stage III checkpoint was scored with a 131k-token input window. For each trait group and distance bin, we reported both AUROC values and their relative change, (AUROC_131*k*_ − AUROC_8*k*_)/AUROC_8*k*_. This end-to-end contrast measures the benefit of long-context scaling rather than a fixed-input model difference; per-variant scores and additional 1M-context records are retained in the source tables.

### 4.9. MSA architecture adaptation

Post-training used a modified CENO implementation rather than the original single-stream Nemotron-H inference path. The original backbone treats the input as one causal sequence. For MSA post-training, we adapted this path to represent the input as a sequence of related rows, following the family-level modeling idea of PoET, which separates within-sequence modeling from between-sequence conditioning in an autoregressive model of homologous sequences [15]. In CENO, each nucleotide token was assigned a row identity, and the post-training stack used a layer schedule that switches between row-local processing and cross-row fusion.

In row-local layers, recurrent scans were segmented at row boundaries and attention visibility was restricted to tokens from the same row before the usual causal mask was applied. In fusion layers, the model operated on the packed causal context and could exchange information across the target and homologous rows. This gives CENO a row-local/fusion topology while keeping the pretrained hybrid Mamba–attention–MoE backbone, nucleotide vocabulary and language-modeling head unchanged.

We post-trained the 300M, 600M and 1B CENO backbones from staged checkpoints. Checkpoint names encode model scale, training stage and architecture index. The selected checkpoints used scale-specific interleaved schedules of row-local and fusion layers (Table S4).

### 4.10. MSA-context post-training

We adapted pretrained CENO checkpoints with an alignment-conditioned training signal derived from GPN-MSA [13]. The key transfer was the source of conditioning, not the exact prediction objective. GPN-MSA uses an alignment-conditioned masked nucleotide prediction objective, in which the masked target position can use both flanking sequence and homologous alignment columns. CENO remains a causal autoregressive model, so we converted this idea into next-token prediction on packed real MSA contexts. Each training example consisted of a target reference sequence and homologous rows sampled from the human multispecies alignment store.

Let *x* = (*x*_1_, . . ., *x_T_*) denote the packed MSA sequence after concatenating the target and homologous rows, and let *w_t_* denote the training weight for position *t*. Post-training minimized a weighted causal negative log-likelihood,

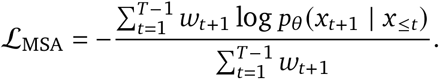

Padding positions and row-boundary transitions were excluded from the loss, so the model was not trained to treat the last nucleotide of one row as the biological predecessor of the first nucleotide of the next. No mask token was inserted, and no bidirectional prediction loss was used. Thus the post-training objective changes the causal sequence likelihood used for downstream variant-effect prediction, but does not introduce a supervised variant classifier or task-specific decision boundary.

The target sequence and homologous rows were concatenated into a single packed sequence before being passed to the model. The row-local/fusion pattern described above interleaves within-row nucleotide modeling with cross-row context exchange in a single causal stack.

### 4.11. Zero-shot variant scoring after post-training

Post-trained CENO models were evaluated with a likelihood-delta interface matching the alternate-versus-reference scoring principle used by GPN-MSA [13]. For each variant, we constructed matched packed contexts for the reference and mutant alleles, keeping the MSA-derived homologous rows fixed across the two scores. Let *C* denote this fixed homologous context and let *y* denote the target row. We scored each target row by its mean causal log-likelihood,

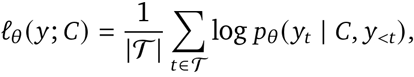

where T contains the scored target-row positions. The variant delta was then

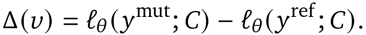

Thus negative Δ(*v*) values indicate that the mutant sequence is less likely than the reference sequence under the same MSA context. Binary classification metrics use −Δ(*v*) as the deleteriousness score, whereas odds-ratio analyses use the signed delta directly with lower scores treated as more deleterious.

### 4.12. Cross-species Fisher odds ratios

For the cross-species enrichment benchmarks, all scores were oriented so that lower values indicate stronger predicted deleteriousness. For CENO this score is the signed likelihood delta Δ(*v*); for conservation baselines with the opposite native direction, scores were negated before thresholding. Let *s_i_* denote the oriented score and *y_i_* ∈ {0, 1} the task label, with *y_i_* = 1 for the positive or rare class and *y_i_* = 0 for the negative or common class. For each model and task, the cutoff was set from the negative class,

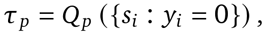

with *p* = 1% in Figure 5f. The low-score tier was defined by *s_i_* ≤ *τ_p_*. We then counted

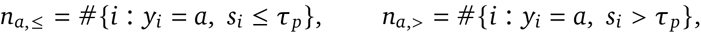

formed the contingency table

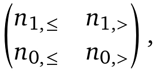

and applied a one-sided Fisher exact test with alternative “greater”. The reported odds ratio was

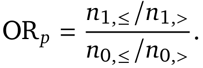

Thus OR*_p_* > 1 indicates enrichment of positive or rare variants in the lowest-scoring tier after controlling the negative-class threshold.

### 4.13. Partial-gene continuation

We randomly selected 100 genes from each of eight species spanning three domains of life: three eukaryotes (*S. cerevisiae*, *D. melanogaster*, *H. sapiens*), three bacteria (*E. coli*, *B. subtilis*, *S. coelicolor*) and two archaea (*S. solfataricus*, *H. salinarum*). For each gene, the first 30%, 50% or 80% of the nucleotide sequence was supplied as a prompt, and the model generated the remaining sequence with vLLM. Recovery rate was computed by comparing the generated continuation with the withheld natural sequence. Figure 4b reports the 80% prompt setting. CENO checkpoints were the 1B, 600M and 300M models after long-context continuation; baselines were Evo 2 7B, Evo 2 7B base, Evo 2 1B base, Evo 1 131k base and Evo 1 8k base. All models were evaluated on the same genes and prompt fractions.

### 4.14. Structure-guided insertion design

#### Design loop

CENO proposed insertion sequences for a target locus; Akita [32] scored candidates and drove an oracle-guided beam search; the surviving candidate set was frozen; and AlphaGenome, which took no part in selection, scored the frozen set across five positional shifts spanning ±8,192 bp. The selection manifest records that the held-out model was not used for selection.

#### Loci, insertion lengths and targets

At 4q25 the insertion is 15,312 bp, the search target is an Akita H1-hESC map and the held-out target is an AlphaGenome H9 ventricular-cardiomyocyte (day 80) map, so evaluation crosses both model and cell type. At LMO2 the insertion is 26,629 bp and both models score GM12878. At IGF2 the insertion is 15,312 bp and both models score HCT116. Displayed contact maps are cropped to 2,048 bp bins, ±164 kb at 4q25 and LMO2 and ±98 kb at IGF2, with a narrow band around the main diagonal masked at plotting time only.

#### Candidate groups

Each boundary-deletion locus was evaluated with matched-random, motif-only and CENO groups; the CENO groups comprise beam-search output plus oracle-ranked and unranked samples drawn without ranking. Group sizes are 160 candidates at 4q25 (five groups of 32) and 136 at LMO2 (four groups of 32 plus eight beam candidates). Beam candidates within a locus share a search lineage; mean pairwise sequence identity is 81% at 4q25 and 92% at LMO2, against 25–28% among the control groups, and effective sample size is correspondingly smaller than the nominal *n*.

#### Recovery at the boundary-deletion loci

Recovery is scaled so that the wild-type contact map is 1 and the deletion map is 0, computed per positional shift and summarized as the group median.

#### Promoter–super-enhancer recovery at IGF2

Because the duplication creates a neo-domain rather than removing a boundary, we scored the promoter–super-enhancer contact directly, with one definition for both models, (*C*_neutral_ − *C*_candidate_)/(*C*_neutral_ − *C*_WT_), where *C*_neutral_ is a length-matched neutral insertion rather than the disease allele, so that recovery discounts the effect of inserting any sequence of that length. AlphaGenome values are the mean over the five positional shifts. The Akita value derives from a single evaluation, so the two models are not matched on window count.

#### 4.15. Cell-type-specific enhancer prediction

The cell-type-specific cis-regulatory element (CRE) prediction and design workflow consists of two components. The first component is a sequence-to-accessibility oracle that predicts chromatin accessibility across mouse cortex subclasses from DNA sequence. The second component is a conditional enhancer generator that uses the oracle as a sequence-level scoring function during supervised finetuning and reinforcement learning. The oracle and the generator share the same pretrained DNA language-model backbone but are trained for different objectives. The same frozen oracle is used consistently to filter supervised fine-tuning examples, to define the reinforcement-learning reward, and to score generated sequences, so that data selection, optimization and evaluation share one accessibility model.

#### Cell-type-specific accessibility oracle

The oracle takes a 2,114-bp DNA sequence as input and predicts accessibility scores across 19 mouse cortex subclasses. The model uses the 600M CENO DNA language-model checkpoint as an unfrozen backbone. DNA sequences are tokenized with the byte-level DNA tokenizer and passed through the backbone to obtain last-layer hidden states. For an input batch, the backbone produces hidden representations with shape [*B*, 2114, 1024]. These representations are then used as input channels for a dilated convolutional neural network head, followed by pooling and a linear output layer that produces a [*B*, 19] vector of subclass-resolved accessibility predictions.

The oracle was trained using a mouse cortex ATAC-seq dataset from the BICCN competition. Candidate CRE regions were represented by fixed 2,114-bp genomic windows centered on the corresponding peak regions. The target matrix contains accessibility values for the 19 mouse cortex subclasses. To reduce sequence leakage between training and evaluation, we used a chromosome-level split: chromosomes 8 and 10 were used for validation, chromosomes 9 and 18 were used for testing, and the remaining chromosomes were used for training.

Oracle training was performed in two stages. In the first stage, the model was trained on all consensus peaks to learn a general sequence-to-accessibility mapping. In the second stage, the model was further trained on cell-type-specific peak regions, which are preferentially accessible in particular mouse cortex subclasses. This two-stage strategy was used to retain broad accessibility prediction while increasing emphasis on cell-type-specific signals.

The training objective combines a pattern-matching term and a magnitude-matching term:

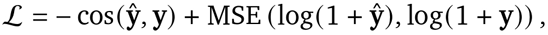

where **y**^ is the predicted accessibility vector and **y** is the target accessibility vector. The cosine term compares the relative accessibility pattern across cell types, whereas the log-MSE term penalizes differences in signal magnitude after log transformation. Training used Adam optimization, learningrate reduction on validation plateau, early stopping, reverse-complement augmentation and small stochastic sequence shifts.

#### In silico oracle evaluation

The oracle was evaluated on chromosome-held-out test regions, including both consensus peaks and cell-type-specific peaks. Prediction accuracy was quantified by comparing predicted and observed accessibility values across test regions and cell types using Pearson correlation, Spearman correlation and related regression metrics.

#### In vivo oracle evaluation

We further evaluated the oracle using a retrospective set of AAV-tested cortical enhancers. The enhancer set contains mouse-derived candidates in mm10 coordinates and human-derived candidates with provided mm10 orthologous liftover coordinates. Mouse-derived enhancers were used with their original mm10 coordinates. Human-derived enhancers were mapped to mm10 using the provided orthologous coordinates, and entries without an orthologous mapping were excluded.

Because the in vivo enhancer annotations and the oracle output use different cell-type taxonomies, the reported in vivo categories were mapped to the 19 oracle subclasses before evaluation. For each enhancer, oracle scores were computed across the mapped subclasses. Candidate enhancers were ranked by the oracle score for the corresponding target class.

Retrospective in vivo evaluation used complementary readouts. Epifluorescence labels were converted into a rank-based metric using the reported enhancer category and signal strength. SSv4 single-cell quantification was used as an additional metric by weighting enhancer categories with the measured cell-type fraction. We also computed precision, recall and F1 in a multi-label setting, using the reported active cell type or cell types as ground truth. In addition, normalized enrichment scores (NESs) were calculated for enhancer categories in the ranked list to quantify category enrichment among high-ranked candidates.

#### Conditional supervised fine-tuning for CRE generation

The conditional enhancer generator was initialized from the 600M CENO DNA language-model checkpoint. Supervised fine-tuning was used to adapt the model from general DNA sequence modeling to conditional CRE generation. Cell-type conditions were represented as single raw-text prefix tokens. Because the tokenizer operates at the byte level, cell-type labels for the four design targets, Astro, Oligo, Pvalb and Sst, were mapped to byte slots that are unused by the nucleotide alphabet, so they were encoded directly without modifying the vocabulary or the embedding table.

Each supervised fine-tuning example consisted of a single cell-type prefix token followed by a target CRE sequence, denoted schematically as

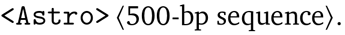

The supervised objective was autoregressive next-token prediction over the CRE sequence conditioned on the prefix. Under this formulation, the model learns the conditional sequence distribution

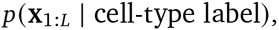

where **x**_1:*L*_ is a 500-bp CRE sequence.

The supervised fine-tuning dataset was constructed from mouse cortex candidate CREs. For each candidate region, a 2,114-bp window was extracted from the mm10 reference genome, and the central 500 bp of this window was used as the generation target; regions whose central 500 bp contained characters outside A/C/G/T were removed. For each of the four design targets, candidate CREs were scored with the frozen oracle, and a target-specific MinGap score was computed over the four design classes:

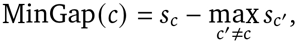

where *s_c_* is the oracle score for the target cell type and the maximum is taken over the other three design classes. CREs with MinGap ≥ 5 were retained as high-specificity training examples, and a fixed number of top-scoring regions was retained per cell type (up to 8,000 for training and 1,000 for validation), with training and validation subsets selected separately per class. Reverse-complement augmentation was applied during data preparation.

To emphasize the most cell-type-selective examples without discarding distributional diversity, the per-token cross-entropy loss was weighted by a monotone function of the example MinGap,

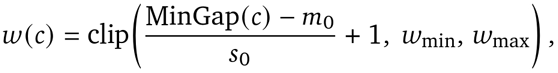

with *m*_0_ = 10, *s*_0_ = 10, *w*_min_ = 0.5 and *w*_max_ = 2.0, so that an example at MinGap = 10 receives unit weight, high-specificity examples are up-weighted up to 2×, and the least specific retained examples are mildly down-weighted. Supervised fine-tuning used the AdamW optimizer with a cosine schedule for six epochs.

#### Oracle-guided reinforcement learning

After supervised fine-tuning, the generator was further optimized with oracle-guided reinforcement learning, initialized from the SFT checkpoint. Optimization used group relative policy optimization (GRPO) [50] with the decoupled-clip and dynamic-sampling policy optimization (DAPO) objective [51]. During this stage, each prompt was the single cell-type prefix token, and the model autoregressively generated a 500-bp DNA sequence. Generated sequences were filtered to retain valid DNA sequences, center-padded with N to the 2,114-bp oracle input length, and scored by the frozen oracle.

The base reward for a generated sequence **x** and target cell type *c* was the oracle MinGap over the four design classes,

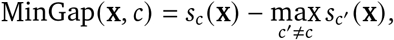

where *s_c_* (**x**) is the oracle-predicted activity for the target cell type and the maximum is taken over the other three design classes. To concentrate the learning signal on high-specificity completions, we used a tail-only shaping of this quantity,

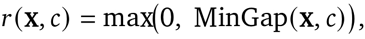

and assigned a fixed negative reward to sequences that failed the validity filter. During GRPO training, multiple completions were sampled from the same cell-type prompt and optimized using relative reward differences within the sampled group. Training used 32 completions per prompt, sampling temperature 0.9, a KL-regularization coefficient of 0.03 toward the supervised fine-tuned reference, group-level reward scaling, and was run for 300 optimization steps; no demonstration sequences were injected into the sampled groups. The final generator was selected from this run. Supervised fine-tuning and reinforcement-learning hyperparameters are summarized in Supplementary Table S7.

## Data and code availability

### Data availability

The datasets used to train and evaluate CENO were obtained from the public resources and benchmarks described in the Methods and cited throughout the report. These third-party datasets remain available from their original providers under the corresponding access conditions and licences.

### Code availability

Code for loading CENO checkpoints, long-context sequence generation and MSA-based variant-effect prediction is publicly available at https://github.com/CladeTeam/CENO. Pretrained CENO and MSA-post-trained CENO-P checkpoints are available from the CENO collection on Hugging Face: https://huggingface.co/collections/CladeTeam/ceno.

## Supplementary Material

### A. Supplementary Figures

**Supplementary Fig. S1.**
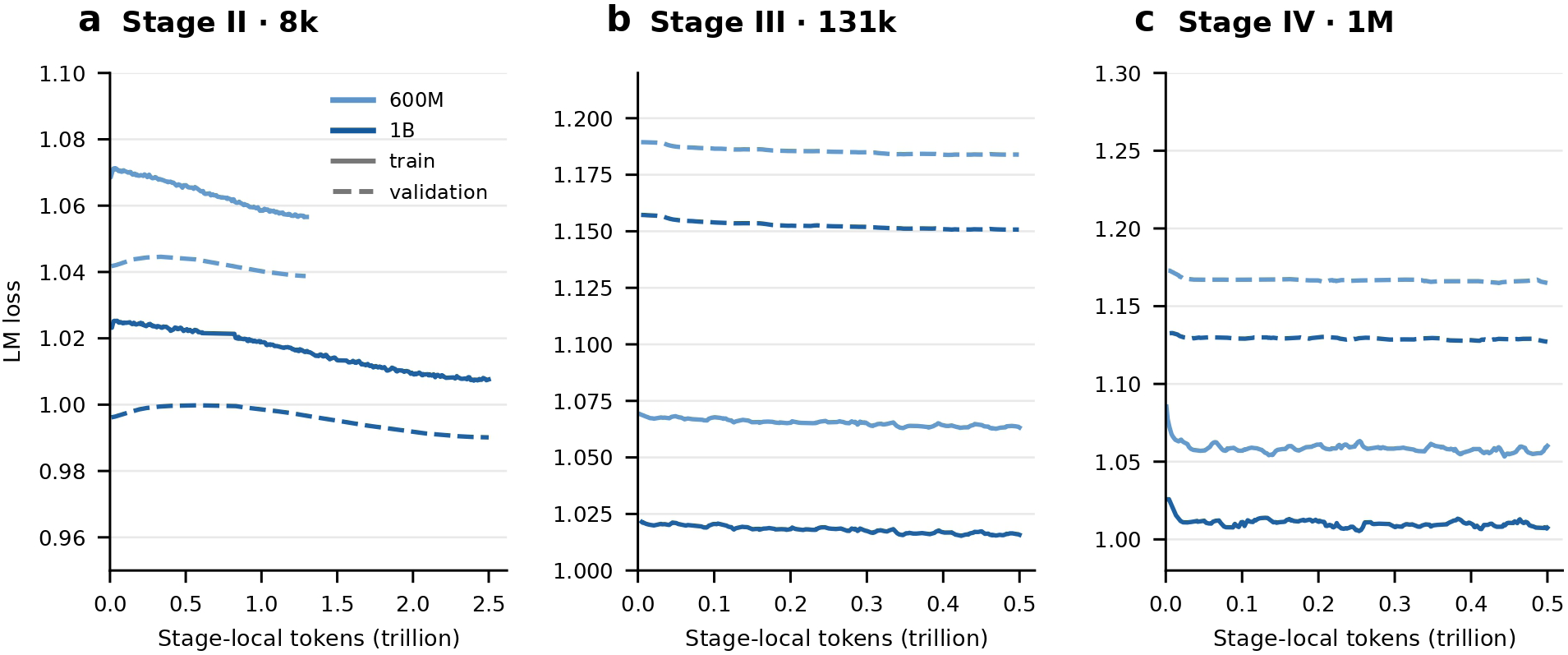
Per-stage training and validation loss for the continued-pretraining curriculum. Training (solid) and validation (dashed) language-modeling loss versus stage-local tokens for the 600M and 1B CENO models during **a,** Stage II (8k-token context), **b,** Stage III (131k-token context) and **c,** Stage IV (1M-token context). Filled circles mark stage-end validation values (annotated). Curves are lightly denoised (robust outlier rejection followed by median/EWM smoothing); context length and data mixture differ across stages.

**Supplementary Fig. S2.**
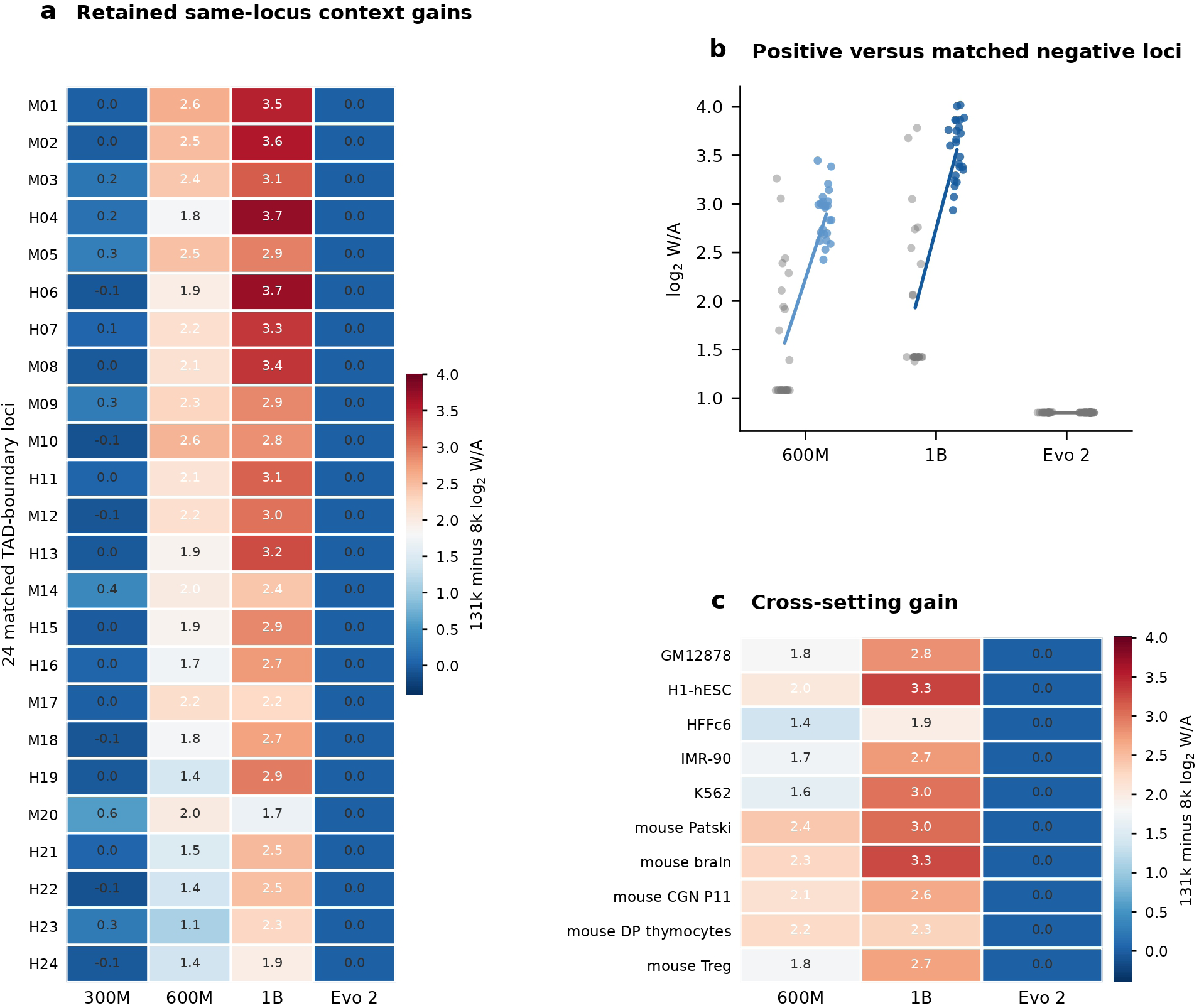
TAD attention samples and cross-setting gains. **a,** Same-locus 131k-minus-8k log_2_ within/across-TAD attention (W/A) gains for 24 retained matched TAD-boundary loci, ranked by CENO gain (rows labeled H, human; M, mouse). **b,** Per-locus log_2_ W/A for matched negative controls (left, gray) versus positive boundaries (right, colored) in the 600M, 1B and Evo 2 7B 131k settings; connecting lines join the group means. **c,** Cross-setting 131k-minus-8k log_2_ W/A gains across five human and five mouse cell or tissue settings. The color scale in **a** and **c** is shared.

**Supplementary Fig. S3.**
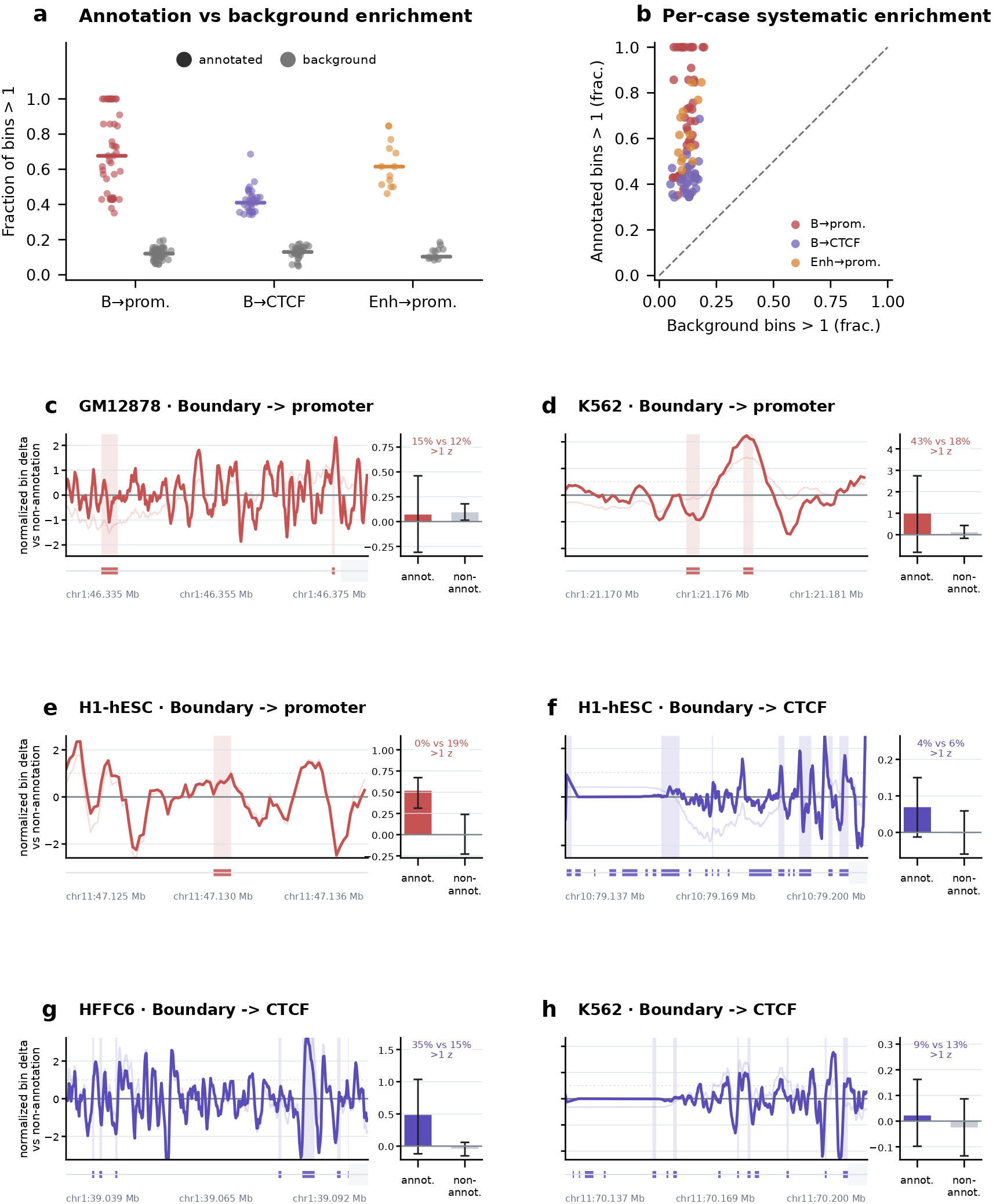
Annotation-linked attention: overall statistics and representative examples. **a,** Fraction of bins exceeding one normalized attention unit for annotated (colored) versus background (gray) bins across all successful cases in the boundary→promoter, boundary→CTCF and enhancer→promoter relationships; bars denote medians. **b,** Per-case comparison of the same fractions (background, *x*; annotated, *y*); all cases lie above the equality line, indicating a systematic rather than cherry-picked effect. **c–h,** Six representative CENO 1B 131k query-to-annotation profiles that are typical rather than top-ranked (three boundary→promoter, three boundary→CTCF; GM12878, K562, H1-hESC, HFFc6 and IMR-90). Traces show the normalized attention delta of the boundary query relative to non-annotation bins along the locus; shaded spans mark annotation bins, with the genomic coordinate track below. Insets give the mean normalized attention in annotated versus non-annotated bins (error bars, bootstrap 95% CI) and the percentage of bins above one *z*-unit.

**Supplementary Fig. S4.**
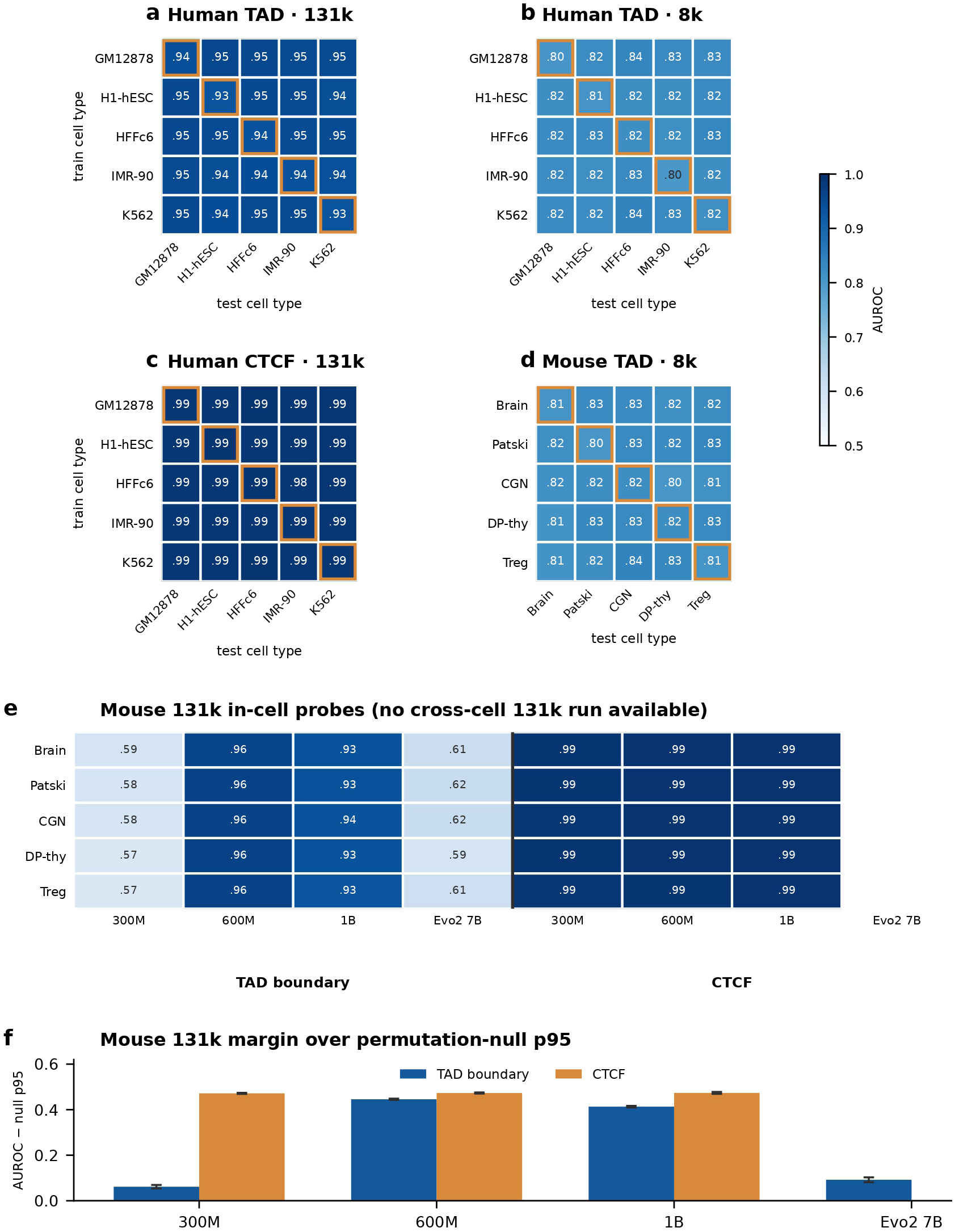
Frozen-state cross-cell-type TAD-boundary and CTCF probe transfer. **a–d,** Train × test AUROC matrices for frozen-representation linear probes evaluated at the layer maximizing in-cell AUROC; orange boxes mark the in-cell diagonal (train=test). **a,** Human TAD boundary, 131k; **b,** human TAD boundary, 8k; **c,** human CTCF, 131k; **d,** mouse TAD boundary, 8k, the only context and probe with a mouse cross-cell-type run. **e,** Mouse 131k in-cell selected-layer AUROC for TAD-boundary and CTCF probes across cell or tissue settings and model scales; no mouse cross-cell-type run exists at 131k (Evo 2 7B CTCF not run). **f,** Mouse 131k selected-layer AUROC margin over the label-shuffle permutation-null p95 (mean ± s.d. across the five settings); all margins are positive.

**Supplementary Fig. S5.**
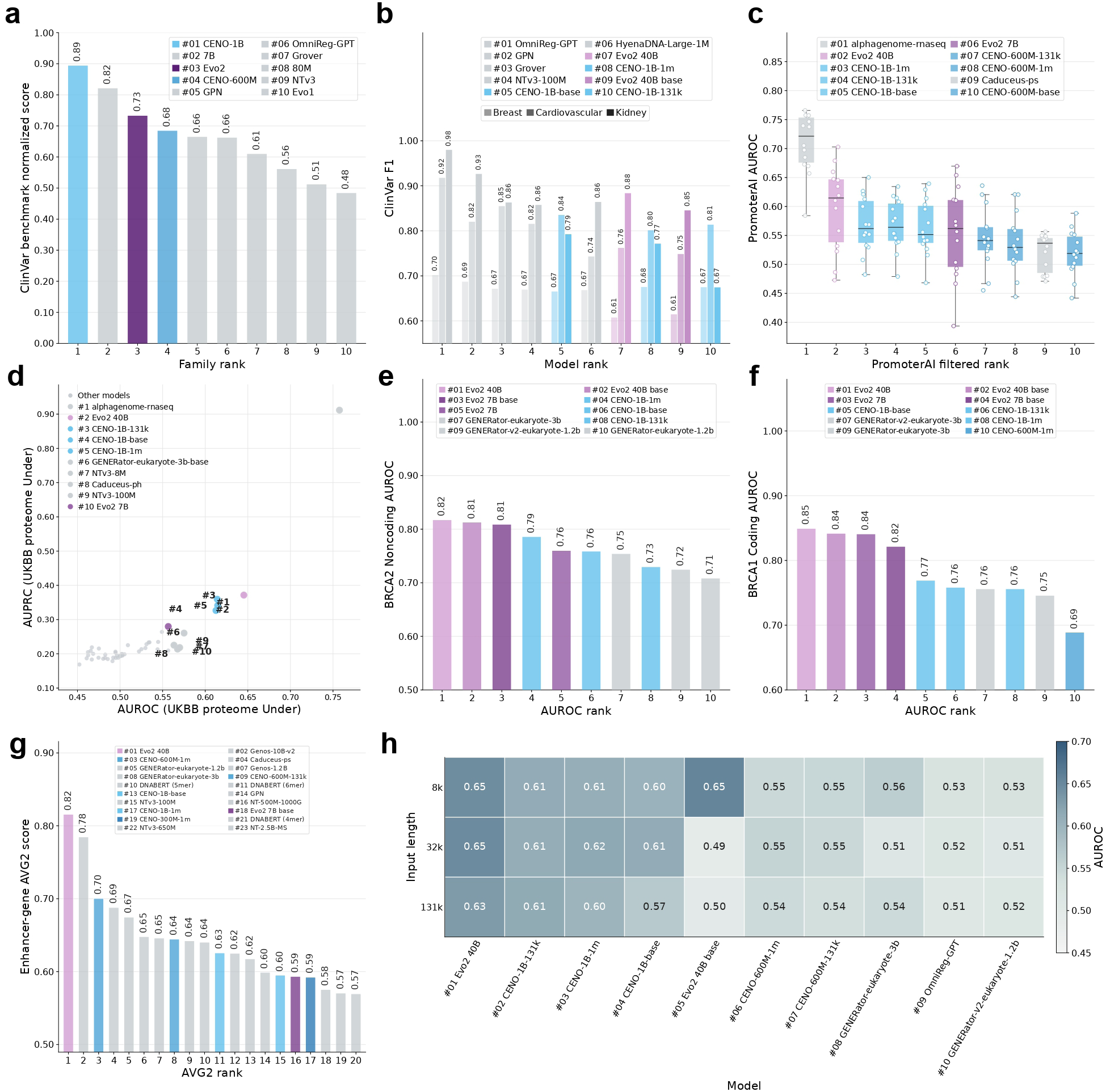
Supplementary task-specific clinical, regulatory and BRCA benchmarks. **a,** ClinVar family-level normalized score, with CENO model sizes shown separately. **b,** ClinVar F1 across clinical groups. **c,** PromoterAI filtered-rank summary across expression-perturbation subtasks. **d,** PromoterAI UKBB proteome under-expression AUROC–AUPRC frontier. **e,** BRCA2 noncoding SNV AUROC. **f,** BRCA1 coding SNV AUROC. **g,** Enhancer-gene AVG2 ranking. **h,** TraitGym AUROC by input length for the top-ranked models.

**Supplementary Fig. S6.**
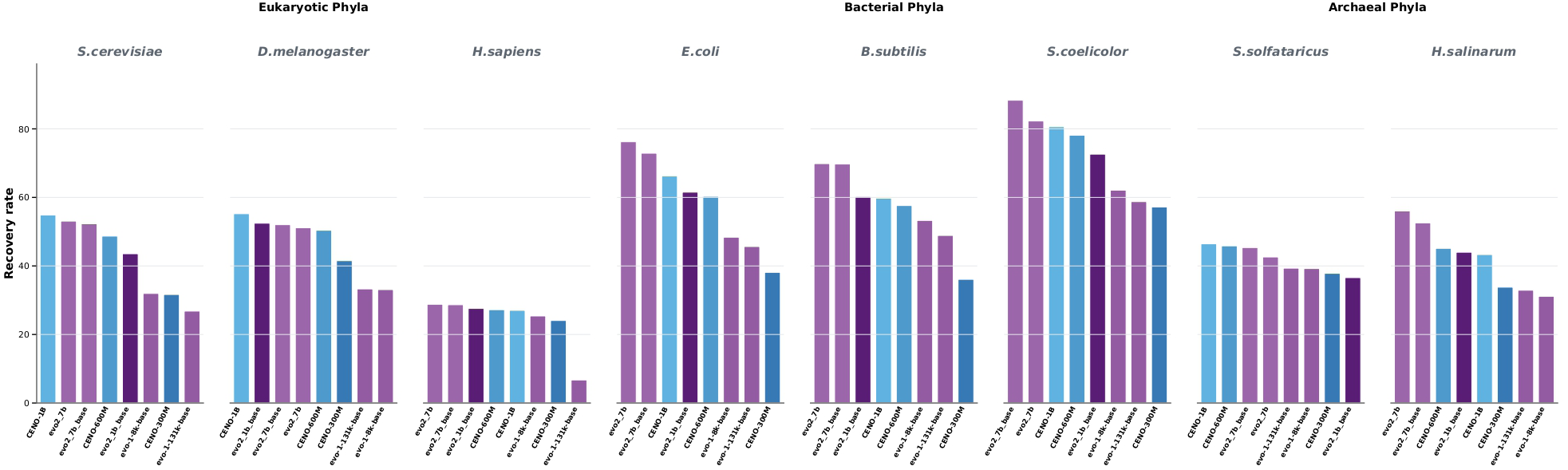
Gene-recovery analysis across species. The analysis used conserved housekeeping genes from diverse eukaryotic, bacterial and archaeal phyla to evaluate sequence recovery and taxonomic coverage across the three domains of life.

**Supplementary Fig. S7.**
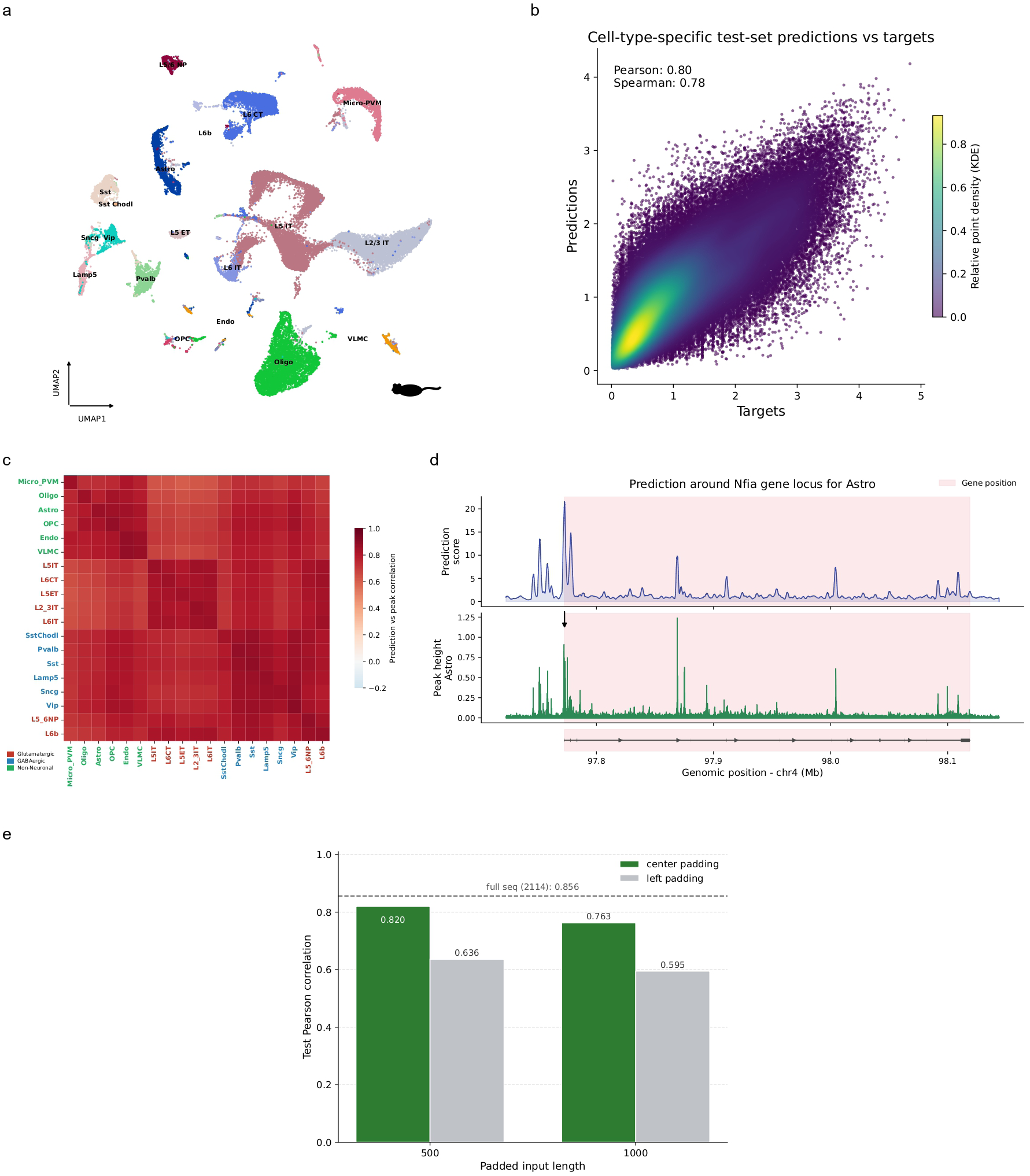
Evaluation of the cell-type-specific accessibility oracle in mouse cortex. **a,** UMAP visualization of mouse motor cortex cells from the BICCN multiome dataset, colored by subclass annotation. **b,** Density scatter plot comparing oracle predictions and observed chromatin accessibility on held-out cell-type-specific peaks. Each point represents one nonzero peak–cell-type pair after log transformation, and color indicates relative point density estimated by KDE. **c,** Cross-cell-type correlation matrix between predicted and observed chromatin accessibility profiles across held-out consensus peaks. Rows and columns correspond to mouse cortex subclasses, with label colors indicating major cell groups. **d,** Locus-level oracle scoring at the astrocyte marker gene *Nfia*. The top track shows the predicted astrocyte accessibility profile generated by sliding-window scoring, and the bottom track shows the observed astrocyte ATAC signal from the matched BigWig track. The shaded region marks the *Nfia* gene body. **e,** Effect of padded input length and padding strategy on oracle performance. Bars show test Pearson correlation for center padding and left padding at 500 bp and 1,000 bp padded input lengths; the dashed line indicates the full 2,114 bp input setting.

**Supplementary Fig. S8.**
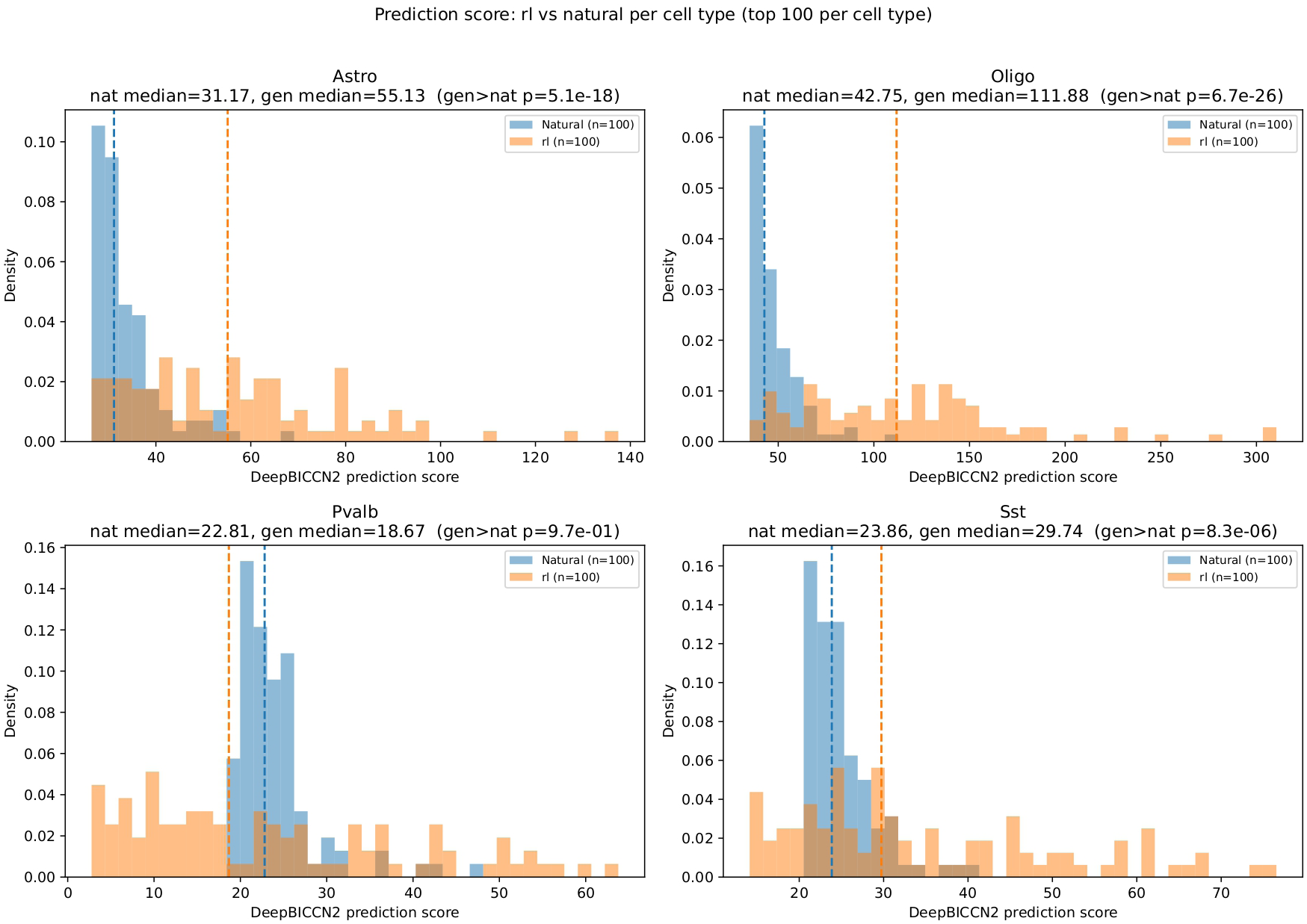
Target-cell oracle score distributions of RL-generated and natural cell-type-specific CREs. Histograms compare on-target oracle prediction scores between RL-generated sequences and natural cell-type-specific CREs for Astro, Oligo, Pvalb and Sst. For each cell type, the top 100 sequences ranked by target-cell score were used for both groups. Dashed lines indicate group medians. Statistical significance was assessed using a one-sided Mann–Whitney U test for higher scores in generated sequences.

**Supplementary Fig. S9.**
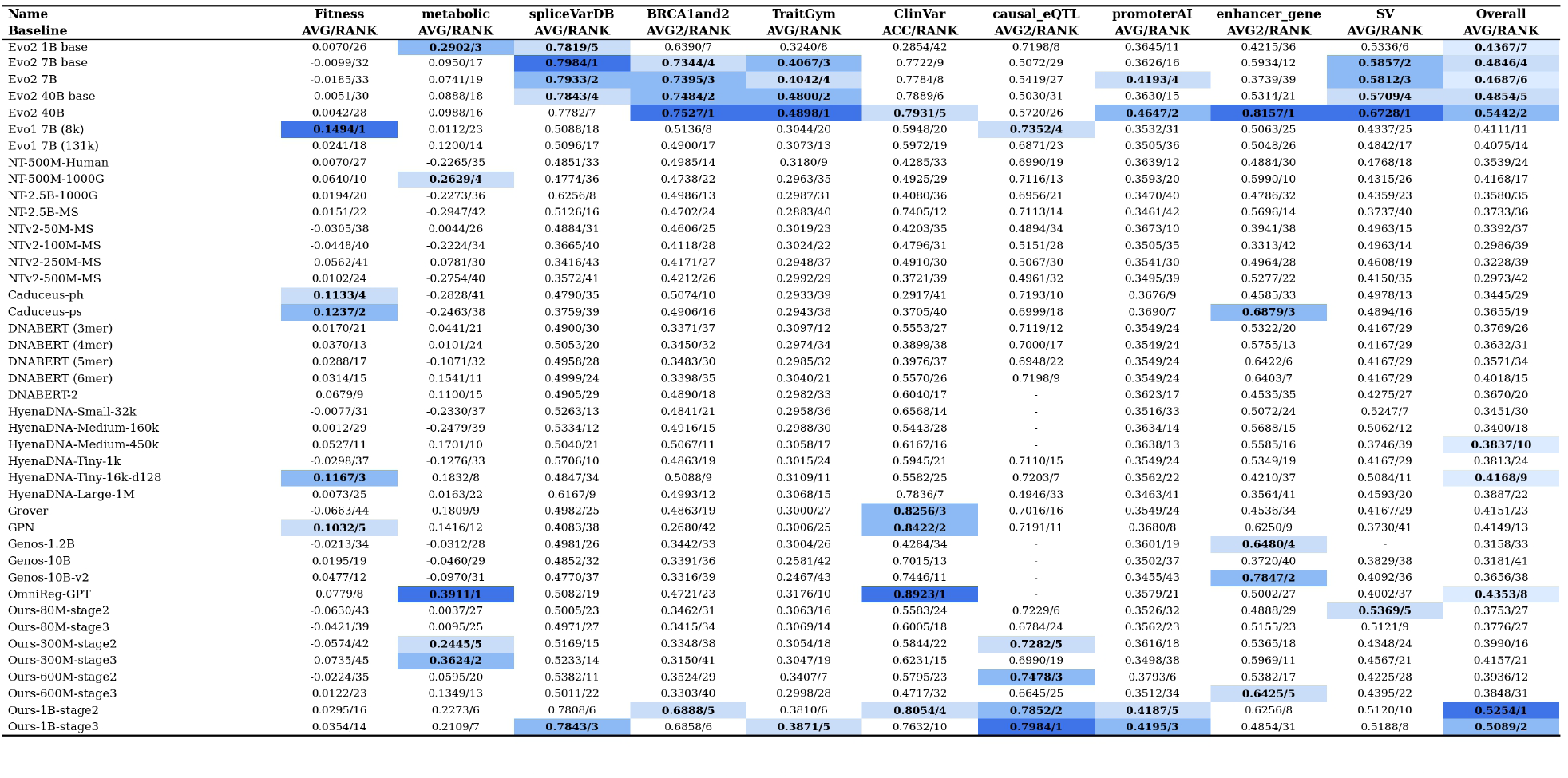
Variant effect prediction benchmark results. Each entry reports the source score and its rank in the form score/rank.

### B. Supplementary Tables

**Supplementary Table S1.** Model configuration and pretraining recipe. Block notation: M, Mamba-2 sequence-mixing block; E, MoE feed-forward block with 8 experts and top-2 routing; *, attention block; -, dense feed-forward block.

| Model | Total | Active | Layers | Hidden | Heads | Max context | Blocks | Layer pattern |
| --- | --- | --- | --- | --- | --- | --- | --- | --- |
| 300M | 300M | 100M | 9 | 1,024 | 8 | 1,048,576 | M4 / E4 / *1 / -0 | MEM*EMEME |
| 600M | 600M | 200M | 20 | 1,024 | 16 | 1,048,576 | M9 / E8 / *2 / -1 | M-MEM*EMEMEM*EMEMEME |
| 1B | 1B | 400M | 38 | 1,024 | 16 | 1,048,576 | M17 / E17 / *4 / -0 | MEMEM*EMEMEM*EMEMEMEMEM*EMEMEM*EMEMEME |

| Model | Stage | Context | Tokens | Samples | GPUs | CP | Micro batch | Global batch | LR range |
| --- | --- | --- | --- | --- | --- | --- | --- | --- | --- |
| 300M | I | 8k | 1.0T | 122.1M | 64 | 1 | 32 | 1,024 | $1 \times 10^{-3}$–$1 \times 10^{-4}$ |
| 300M | II | 8k | 0.5T | 61.0M | 64 | 1 | 32 | 1,024 | $1 \times 10^{-4}$–$1 \times 10^{-5}$ |
| 300M | III | 131k | 500B | 3.8M | 32 | 1 | 1 | 64 | $2 \times 10^{-5}$–$2 \times 10^{-6}$ |
| 300M | IV | 1M | 500B | 0.5M | 32 | 4 | 1 | 8 | $5 \times 10^{-6}$–$5 \times 10^{-7}$ |
| 600M | I | 8k | 2.0T | 244.1M | 32 | 1 | 16 | 1,024 | $8 \times 10^{-4}$–$8 \times 10^{-5}$ |
| 600M | II | 8k | 1.3T | 158.7M | 32 | 1 | 16 | 1,024 | $8 \times 10^{-5}$–$8 \times 10^{-6}$ |
| 600M | III | 131k | 500B | 3.8M | 32 | 2 | 1 | 64 | $1.6 \times 10^{-5}$–$1.6 \times 10^{-6}$ |
| 600M | IV | 1M | 500B | 0.5M | 64 | 8 | 1 | 8 | $4 \times 10^{-6}$–$4 \times 10^{-7}$ |
| 1B | I | 8k | 3.5T | 427.2M | 64 | 1 | 8 | 1,024 | $5 \times 10^{-4}$–$5 \times 10^{-5}$ |
| 1B | II | 8k | 2.5T | 305.2M | 64 | 1 | 8 | 1,024 | $5 \times 10^{-5}$–$5 \times 10^{-6}$ |
| 1B | III | 131k | 500B | 3.8M | 32 | 4 | 1 | 64 | $1 \times 10^{-5}$–$1 \times 10^{-6}$ |
| 1B | IV | 1M | 500B | 0.5M | 64 | 16 | 1 | 8 | $2.5 \times 10^{-6}$–$2.5 \times 10^{-7}$ |

**Supplementary Table S2.** Data mixture weights. Weights for Stage I, Stage II and the long-context continuation stages. Values are percentages of sampled training tokens.

| Data source | Stage I | Stage II | Stage III/IV |
| --- | --- | --- | --- |
| GTDB / IMG prokaryotic genomes | 22.00 | 8.00 | 18.18 |
| Metagenomes | 24.00 | 14.00 | 10.10 |
| Viral genomes | 3.00 | 4.00 | 3.03 |
| Organelle genomes | 2.00 | 2.00 | 1.01 |
| Eukaryotic genic windows | 25.00 | 35.00 | 16.16 |
| mRNA | 10.00 | 15.00 | 11.11 |
| mRNA splice/promoter windows | 8.00 | 14.00 | 11.11 |
| ncRNA | 3.00 | 5.00 | 4.04 |
| Promoter windows | 3.00 | 3.00 | 1.01 |
| NCBI eukaryotic genomes, Animalia | 0.00 | 0.00 | 16.16 |
| NCBI eukaryotic genomes, Plantae | 0.00 | 0.00 | 5.05 |
| NCBI eukaryotic genomes, Fungi | 0.00 | 0.00 | 2.02 |
| NCBI eukaryotic genomes, Protista | 0.00 | 0.00 | 0.51 |
| NCBI eukaryotic genomes, Chromista | 0.00 | 0.00 | 0.51 |
| Grouped prokaryotic/metagenomic/viral/organelle | 51.00 | 28.00 | 32.32 |
| Grouped eukaryotic/transcript/regulatory/complete-genome | 49.00 | 72.00 | 67.68 |

**Supplementary Table S3.**
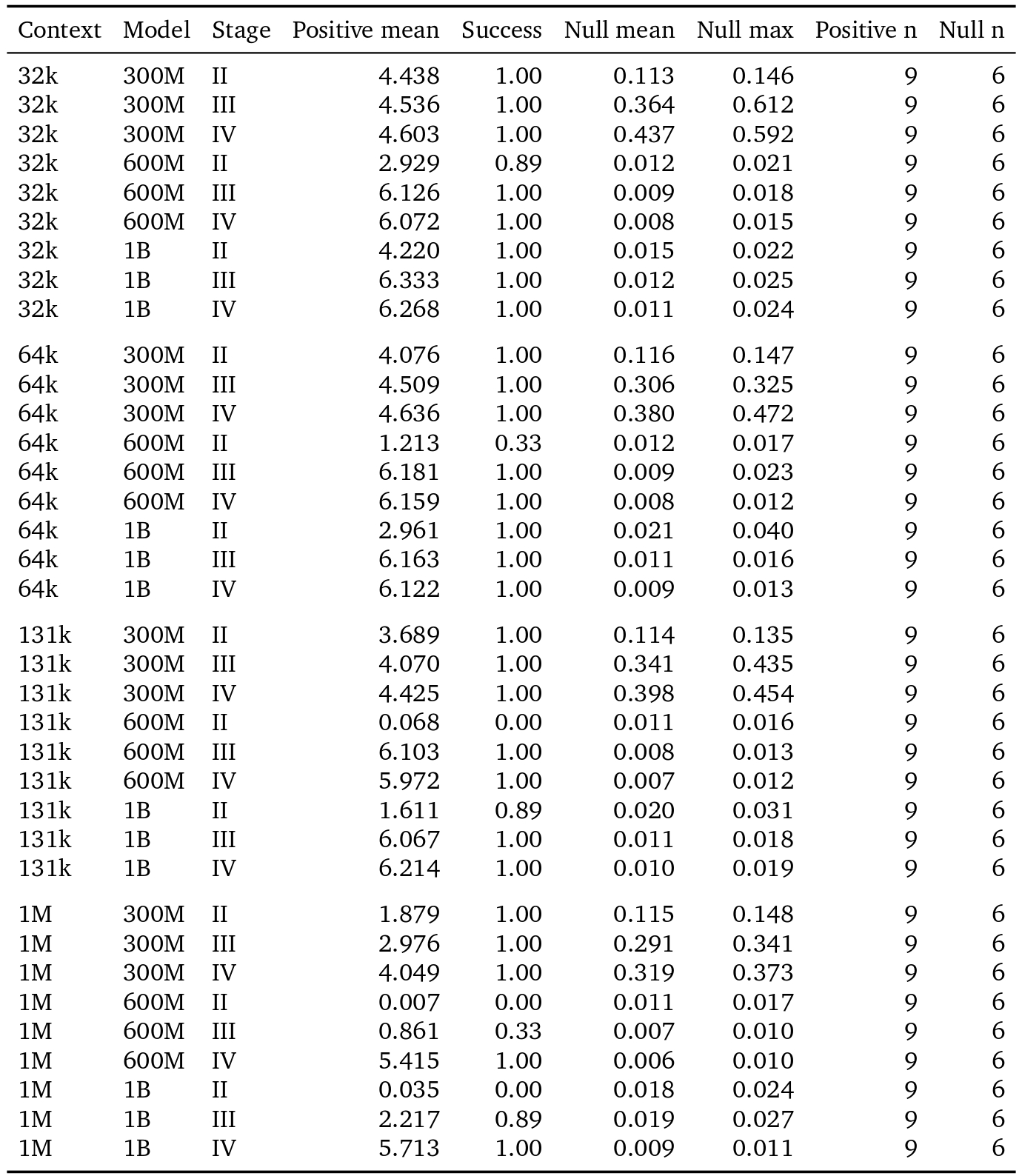
Needle retrieval summary. For each model-stage and test-context setting, positive statistics summarize nine records (three insertion depths by three trials), whereas null statistics summarize six shuffled-needle controls (two insertion depths by three trials). Success denotes the fraction of positive trials exceeding the prespecified retrieval-score threshold of 0.8.

**Supplementary Table S4.**
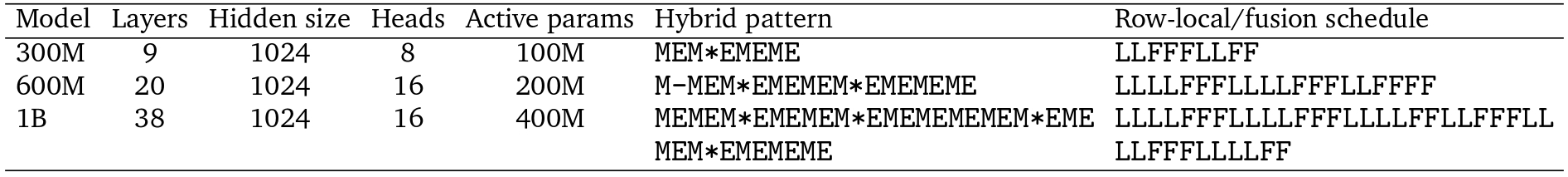
CENO post-training architecture schedules. Hybrid pattern symbols match Table S1. In the row-local/fusion schedule, L denotes a row-local layer that restricts context to the same MSA row, and F denotes a fusion layer that uses the packed causal context across rows.

**Supplementary Table S5.**
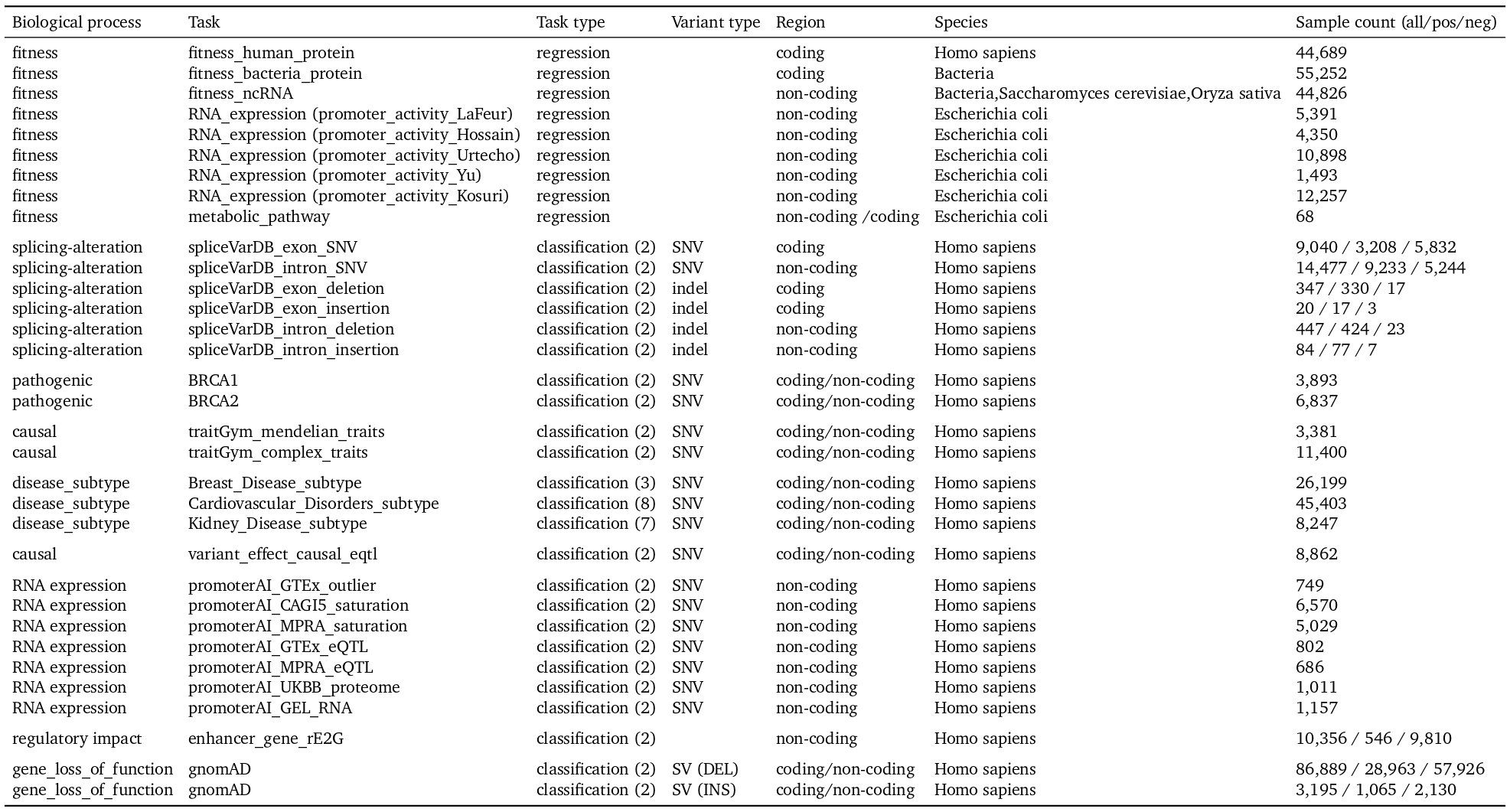
Variant benchmark information. Benchmark tasks are organized by biological process, task type, variant type, genomic region and species. Sample counts are reported as all/positive/negative when class counts are available.

| Biological process | Task | Task type | Variant type | Region | Species | Sample count (all/pos/neg) |
| --- | --- | --- | --- | --- | --- | --- |
| fitness | fitness_human_protein | regression |  | coding | Homo sapiens | 44,689 |
| fitness | fitness_bacteria_protein | regression |  | coding | Bacteria | 55,252 |
| fitness | fitness_ncRNA | regression |  | non-coding | Bacteria,Saccharomyces cerevisiae,Oryza sativa | 44,826 |
| fitness | RNA_expression (promoter_activity_LaFeur) | regression |  | non-coding | Escherichia coli | 5,391 |
| fitness | RNA_expression (promoter_activity_Hossain) | regression |  | non-coding | Escherichia coli | 4,350 |
| fitness | RNA_expression (promoter_activity_Urtecho) | regression |  | non-coding | Escherichia coli | 10,898 |
| fitness | RNA_expression (promoter_activity_Yu) | regression |  | non-coding | Escherichia coli | 1,493 |
| fitness | RNA_expression (promoter_activity_Kosuri) | regression |  | non-coding | Escherichia coli | 12,257 |
| fitness | metabolic_pathway | regression |  | non-coding / coding | Escherichia coli | 68 |
| splicing-alteration | spliceVarDB_exon_SNV | classification (2) | SNV | coding | Homo sapiens | 9,040 / 3,208 / 5,832 |
| splicing-alteration | spliceVarDB_intron_SNV | classification (2) | SNV | non-coding | Homo sapiens | 14,477 / 9,233 / 5,244 |
| splicing-alteration | spliceVarDB_exon_deletion | classification (2) | indel | coding | Homo sapiens | 347 / 330 / 17 |
| splicing-alteration | spliceVarDB_exon_insertion | classification (2) | indel | coding | Homo sapiens | 20 / 17 / 3 |
| splicing-alteration | spliceVarDB_intron_deletion | classification (2) | indel | non-coding | Homo sapiens | 447 / 424 / 23 |
| splicing-alteration | spliceVarDB_intron_insertion | classification (2) | indel | non-coding | Homo sapiens | 84 / 77 / 7 |
| pathogenic | BRCA1 | classification (2) | SNV | coding/non-coding | Homo sapiens | 3,893 |
| pathogenic | BRCA2 | classification (2) | SNV | coding/non-coding | Homo sapiens | 6,837 |
| causal | traitGym_mendelian_traits | classification (2) | SNV | coding/non-coding | Homo sapiens | 3,381 |
| causal | traitGym_complex_traits | classification (2) | SNV | coding/non-coding | Homo sapiens | 11,400 |
| disease_subtype | Breast_Disease_subtype | classification (3) | SNV | coding/non-coding | Homo sapiens | 26,199 |
| disease_subtype | Cardiovascular_Disorders_subtype | classification (8) | SNV | coding/non-coding | Homo sapiens | 45,403 |
| disease_subtype | Kidney_Disease_subtype | classification (7) | SNV | coding/non-coding | Homo sapiens | 8,247 |
| causal | variant_effect_causal_eqtl | classification (2) | SNV | coding/non-coding | Homo sapiens | 8,862 |
| RNA expression | promoterAI_GTEx_outlier | classification (2) | SNV | non-coding | Homo sapiens | 749 |
| RNA expression | promoterAI_CAGI5_saturation | classification (2) | SNV | non-coding | Homo sapiens | 6,570 |
| RNA expression | promoterAI_MPRA_saturation | classification (2) | SNV | non-coding | Homo sapiens | 5,029 |
| RNA expression | promoterAI_GTEx_eQTL | classification (2) | SNV | non-coding | Homo sapiens | 802 |
| RNA expression | promoterAI_MPRA_eQTL | classification (2) | SNV | non-coding | Homo sapiens | 686 |
| RNA expression | promoterAI_UKBB_proteome | classification (2) | SNV | non-coding | Homo sapiens | 1,011 |
| RNA expression | promoterAI_GEL_RNA | classification (2) | SNV | non-coding | Homo sapiens | 1,157 |
| regulatory impact | enhancer_gene_rE2G | classification (2) |  | non-coding | Homo sapiens | 10,356 / 546 / 9,810 |
| gene_loss_of_function | gnomAD | classification (2) | SV (DEL) | coding/non-coding | Homo sapiens | 86,889 / 28,963 / 57,926 |
| gene_loss_of_function | gnomAD | classification (2) | SV (INS) | coding/non-coding | Homo sapiens | 3,195 / 1,065 / 2,130 |

**Supplementary Table S6.** GeneBench recovery rates for overall and house-keeping genes. Values are mean recovery rates at prompt proportion 0.8 and are reported to four decimal places. Overall entries below 10 are shown as unavailable, matching the source plotting rule. For house-keeping genes, CENO results use top-*p* = 0.8; the Evo source files do not expose a top-*p* field. The house-keeping CENO-300M result uses 300M_A100M_stage4_1m_c3_corrected as a proxy because the exact 300M_A100M_stage3_32k checkpoint is absent from the house-keeping results.

| Model | Eukaryotic phyla |  |  | Bacterial phyla |  |  | Archaeal phyla |  |
| --- | --- | --- | --- | --- | --- | --- | --- | --- |
|  | *S. cerevisiae* | *D. melanogaster* | *H. sapiens* | *E. coli* | *B. subtilis* | *S. coelicolor* | *S. solfataricus* | *H. salinarum* |
| **Overall genes** |  |  |  |  |  |  |  |  |
| CENO-300M | 33.9316 | 29.6442 | 29.0910 | 31.1285 | 33.8771 | 40.4032 | 37.5135 | 37.3820 |
| CENO-600M | 39.1846 | 31.6312 | 33.2939 | 44.3932 | 43.5491 | 50.3933 | 40.5506 | 41.4166 |
| CENO-1B | 42.9717 | 35.7403 | 34.9017 | 53.0278 | 45.9558 | 55.3328 | 41.1605 | 43.8755 |
| evo2_1b_base | 32.2276 | 30.2824 | 30.5315 | 47.9408 | 37.6592 | 48.0515 | 32.7150 | 37.0400 |
| evo2_7b_base | 35.9555 | 32.1896 | 30.0951 | 61.2794 | 44.5505 | 58.0307 | 35.8889 | 43.6873 |
| evo2_7b | 36.9694 | 31.8500 | 30.9437 | 65.2630 | 45.6764 | 59.4867 | 35.0630 | 44.5220 |
| evo-1-8k-base | 28.7700 | 28.2007 | – | 35.5491 | 34.8927 | 40.6562 | 32.1771 | 30.4542 |
| evo-1-131k-base | – | 28.4273 | 23.0750 | 35.7043 | 34.7181 | 40.7437 | 31.7141 | 29.8060 |
| **House-keeping genes** |  |  |  |  |  |  |  |  |
| CENO-300M | 31.6604 | 41.5211 | 24.0690 | 38.1189 | 36.0860 | 57.1978 | 37.8237 | 33.7981 |
| CENO-600M | 48.6977 | 50.3925 | 27.2257 | 60.3060 | 57.6117 | 78.1280 | 45.8513 | 45.1247 |
| CENO-1B | 54.8436 | 55.2305 | 27.0263 | 66.2239 | 59.7235 | 80.6081 | 46.4494 | 43.3100 |
| evo2_1b_base | 43.5543 | 52.4613 | 27.5833 | 61.5198 | 60.2469 | 72.6148 | 36.5849 | 43.9949 |
| evo2_7b_base | 52.2830 | 52.0291 | 28.6734 | 72.8877 | 69.7262 | 88.3447 | 45.3591 | 52.5624 |
| evo2_7b | 53.0759 | 51.1342 | 28.8052 | 76.2302 | 69.8433 | 82.2873 | 42.6077 | 56.0407 |
| evo-1-8k-base | 31.9632 | 33.0835 | 25.3951 | 48.3786 | 53.2724 | 62.1117 | 39.2162 | 31.1364 |
| evo-1-131k-base | 26.8170 | 33.2878 | 6.7263 | 45.6498 | 48.8879 | 58.7515 | 39.3396 | 32.9022 |

**Supplementary Table S7.** Cell-type-specific CRE supervised fine-tuning and reinforcement-learning hyperparameters. Settings for the conditional CRE generator built on the 600M CENO backbone. The MinGap contrast is computed over the four design classes (Astro, Oligo, Pvalb, Sst).

| Stage | Setting | Value |
| --- | --- | --- |
| Shared | Backbone | 600M CENO (Stage II) |
| Shared | Conditioning | single raw-text cell-type prefix token |
| Shared | Generation target length | 500 bp (center crop of 2,114-bp window) |
| Shared | Oracle input length | 2,114 bp (center-padded with N) |
| Shared | MinGap contrast classes | Astro, Oligo, Pvalb, Sst |
| SFT | MinGap filter | $\geq 5$ |
| SFT | Per-cell top-$K$ (train / val) | 8,000 / 1,000 |
| SFT | Loss | weighted autoregressive cross-entropy |
| SFT | MinGap weight ($m_0, s_0, w_{\min}, w_{\max}$) | (10, 10, 0.5, 2.0) |
| SFT | Optimizer / schedule | AdamW / cosine |
| SFT | Learning rate | $5 \times 10^{-5}$ |
| SFT | Epochs | 6 |
| SFT | Augmentation | reverse complement |
| RL | Algorithm | GRPO (DAPO objective) |
| RL | Initialization | SFT checkpoint |
| RL | Reward | tail-only oracle MinGap, $\max(0, \cdot)$ |
| RL | Invalid-sequence penalty | applied |
| RL | Completions per prompt | 32 |
| RL | Sampling temperature | 0.9 |
| RL | KL coefficient | 0.03 |
| RL | Reward scaling | group-level |
| RL | Optimization steps | 300 |

